# Predicting Neural Activity from Connectome Embedding Spaces

**DOI:** 10.1101/2025.05.09.653224

**Authors:** Zihan Zhang, Huanqiu Zhang, Stefan Mihalas

**Affiliations:** Center for Data-Driven Discovery in Biology, Allen Institute, Seattle, WA 98109, USA; Data Science Institute & Department of Neurobiology, The University of Chicago, Chicago, IL 60637, USA

## Abstract

Understanding how structured patterns of neural activity emerge from the underlying connectivity is fundamental to elucidating brain function. While cortical connectomes are intrinsically high-dimensional, population activity typically resides in a much lower-dimensional subspace. Consequently, only a small fraction of the information encoded in the connectome appears relevant for shaping activity. Can we identify low-dimensional features of the connectome features be that reliably predict neural activity? Leveraging the MICrONS dataset, which combines millimeter-scale, nanometer-resolution connectivity with simultaneously recorded in-vivo activity, we demonstrate a statistically significant alignment between morphological and functional similarity, quantified by subspace angles and centered kernel alignment. Topological analyses further reveal that the representation spaces of both the connectome and neural activity share a low-dimensional hyperbolic geometry with exponential scaling. These parallels motivated the hypothesis that embedding anatomical affinities into an appropriate geometric space can isolate the functionally relevant features of the connectome. We therefore applied multidimensional scaling to generate such embeddings and trained a simple linear model to reconstruct neuronal activity. Remarkably, the embedded connectome explained 68% in activity similarity, surpassing models that had direct access to activity similarity itself and outperforming similarly simple models that used the full high-dimensional connectome (56%). Our findings uncover a robust structure-function coupling: geometry-aware dimensionality reduction discards much of the connectome’s microscopic detail yet yields superior predictions of neural activity. This suggests that synaptic wiring implicitly encodes an abstract, low-dimensional organization that underlies the observed low-dimensional cortical activity.

## 1: Introduction

The human brain is a complex multiscale structure, encompassing various molecular, cellular, and neuronal phenomena that constitute the foundation of cognition (1, 2). Experimental evidence estimates that the average human brain contains over 86 billion neurons and approximately 10^14^ synaptic connections (3, 4), while the mouse brain comprises approximately 85 million neurons, forming roughly 100 billion synaptic connections (5). Each synapse, depending on its strength, has been suggested to store up to 4.7 bits of information (6). However, are all these structural details essential to define the circuit activity in the brain? For the first time, based on monumental efforts (7), we now have the capability to compare simplified models that link structure to activity in cortical circuits under varying levels of coarse-graining in connectivity.

While detailed structural properties shape brain activity, the extent to which they are necessary for functional modeling remains an open question. Simple artificial neural networks (ANNs) without bio-realistic constraints may not serve as ideal models, as their material and metabolic limitations, synaptic signals, and anatomical scaffolds are not encoded in the same manner as parameters in ANNs. Consequently, the mechanics and functionalities of the brain may not be fully explained or replicated by the scaling laws employed in artificial systems (8–13). Recent studies have found that, despite the complex and high-dimensional nature of individual neuronal responses, population activity across neurons often unfolds with low-dimensional subspaces, particularly in sensory and motor systems (14–17). Although these studies suggest that, due to its low-dimensional nature, activity contains only a smaller amount of information than neurons can independently encode, no link to its emergence from structure is proposed.

Historically, the nontrivial relationship between structural connectivity (**W**) and functional characteristics (**A**) of biological neuronal networks has been widely recognized, as exemplified by classical Hebbian theory and the three-factor learning rule (18, 19). However, most studies exploring this relationship remain theoretical and lack systematic validation with empirical data. Experimental efforts to establish a direct link between structure and function primarily rely on MRI data, which emphasize macroscopic network-level analyses rather than detailed synaptic-level investigations (20–25). More recently, studies on *C. elegans* and fruit flies (*Drosophila Melanogaster*) have provided detailed connectomic data, enabling finer-grained analyses (26–30). Building on this, (31) treat the Drosophila connectome as a prior, uncovering that key circuits rely on only a handful of neurons and enabling whole-brain casual modeling from sparse data. (32) likewise predict each neuron’s response to dynamic visual stimuli by sharing parameters within cell types. While these connectome-based studies employ state-of-the-art deep learning models to uncover complex high-dimensional relationships between structure and function, our work instead focuses on identifying normative, low-dimensional representations of these relationships. Specifically, we adopt a low-dimensional embedding approach based on vectorized connectome similarity, which has been previously used to predict and explain edge-wise functional connectivity in inter-hemispheric homologous regions (33, 34). From a broader perspective, our study aligns with previous work by advocating for the incorporation of biological details into computational models in a simplified yet principled manner that enhances both performance and metabolic efficiency (31, 32). However, unlike prior studies that emphasize the precise details of the connectome, our approach focuses on a lossy compression of connectomic information, capturing essential structural properties while abstracting away fine-grained complexities.

## 2: Results

### 2.1. Structure-Function Alignment

We use the largest functionally-imaged electron microscopy (EM) dataset currently available, MICrONS, which was generated through dense calcium imaging followed by the reconstruction of a millimeter-scale volume (7, 35, 36). This dataset provides nanometer-resolution details on the spatial locations of each cell’s soma and associated synapses, along with classification labels, postsynaptic density (PSD) sizes, and activity recordings of neurons in the primary visual cortex (V1) as well as the anterolateral (AL), lateromedial (LM), and rostrolateral (RL) higher visual areas. The activity data consists of detrended, deconvolved, and stimulus- and behavior-aligned recordings from excitatory neurons spanning layers 2 to 6. Figure 1A-B provides an illustration of neuronal connectivity within the dataset. In Sections 2.1 and 2.3, because we incorporate both structural and functional information, we restrict analyses to neurons that were imaged, reconstructed, and coregistered – 497 unique cells in total – comprising 35-80 neurons per scan across 12 scans (Section 4.1). In Section 2.2, only the connectome information is required for the geometric analysis; accordingly, the available sample expands to 716. To maintain consistency with prior studies, we adopt the same inclusion criteria and preprocessing procedures as in (7, 36).

**Fig. 1.**
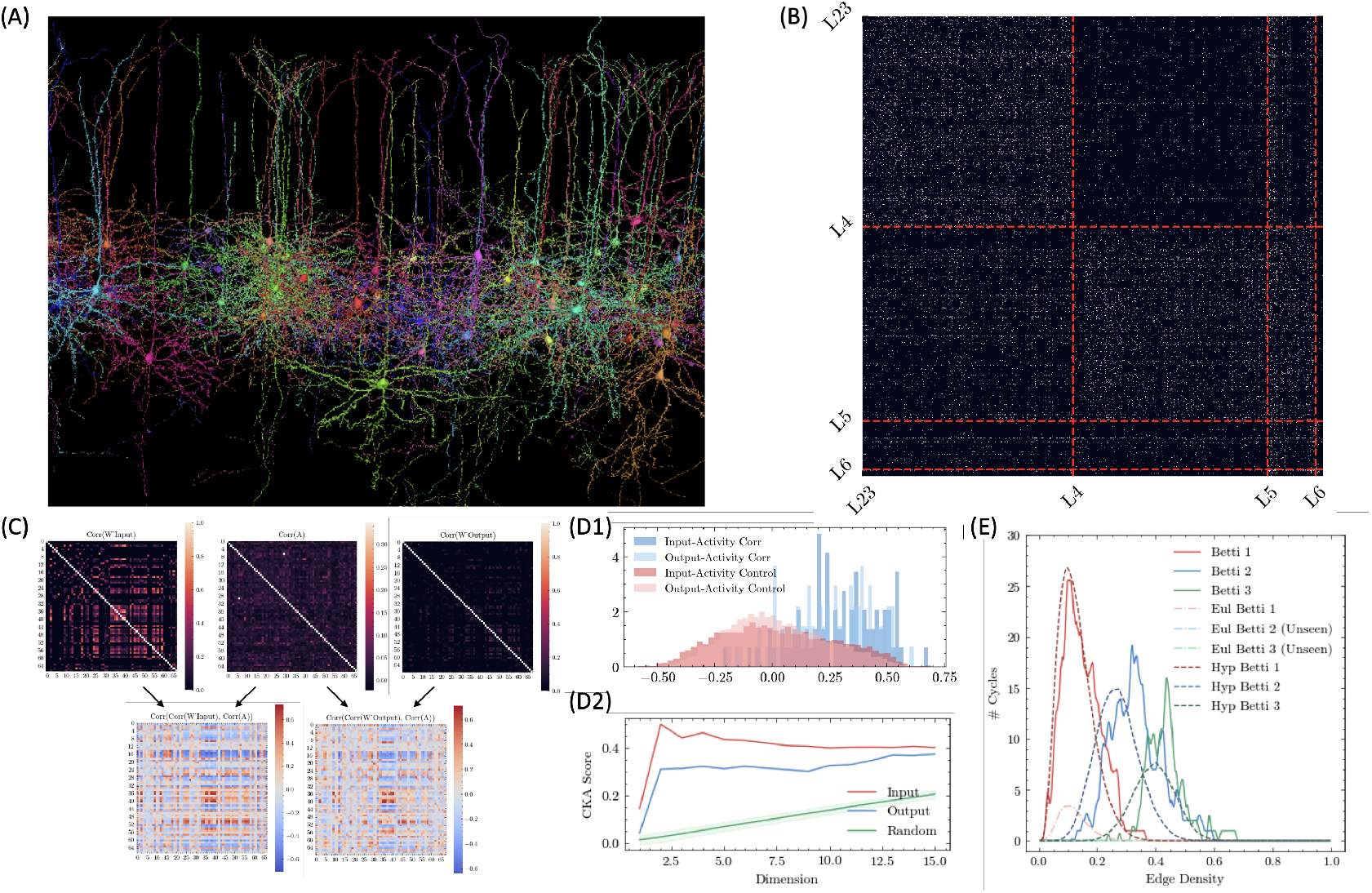
Demonstration of MICrONS Dataset and Structure-Function Alignment: (A): Illustration of 89 proofread layer 4 pyramidal cells from the MICrONS dataset using `neuroglancer`; (B): Binary pairwise connections between fully proofread excitatory neurons (*N* = 716), ordered by layers. The horizontal axis represents postsynaptic neurons, while the vertical axis represents the presynaptic neurons (Section 5.1); (C): Illustration of input connection correlation **C**^in^ (top-left), neuronal activity correlation **C**^act^ (top-middle), output connection correlation **C**^out^ (top-right), **C**^in,act^ (bottom-left, also see Equation (2)), and **C**^out,act^ (bottom-right). For instance, each entry in **C**^in,act^ represents the Pearson correlation between the *i*th row of **C**^in^ and the *j*th row of **C**^act^. This matrix has diagonal elements representing the correlation between connection and activity similarities for each neuron, while the off-diagonal elements serve as controls, relating the functional similarity of one neuron to the connection similarity of another. (D1): Correlation between activity and connection similarity for inputs (dark blue) and outputs (light blue), along their respective controls (dark red and light red). *t*-test *p*-values: *p*_1_ = 2.32 × 10^−22^ (dark blue vs. dark red, testing whether input connections are correlated with activity), *p*_2_ = 2.43 × 10^−16^ (light blue vs. light red, testing whether output connections are correlated with activity), and *p*_3_ = 4.90 × 10^−4^ (dark blue vs. light blue, testing whether input connections exhibit a stronger correlation with activity than output connections). See Figure S2 for results from additional scans; (D2): Centered Kernel Analysis (CKA) of low-rank approximations using Singular Value Decomposition (SVD) for **C**^act^ with **C**^in^ (red), **C**^out^ (blue), and random matrices (green). The *x*-axis represents the inclusion of different numbers of top singular values (i.e. truncation rank) (Section 4.3); (E): Betti curves derived from the activity pairwise correlation matrix **C**^act^ (solid lines) align more closely with points uniformly sampled from two-dimensional hyperbolic space than with those from Euclidean space. Standard deviations are omitted for clarity. Averaging across 100 hyperbolic samples (dark dashed line) at the optimized radius *R*_max_ value (Section 4.2) yields a cumulative absolute pointwise error of 14.67 across the first three Betti curves. The corresponding hyperbolic Betti values, 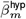 for *m* = 1, 2, 3, are 3.31, 2.46, and 1.21, which closely match the groundtruth Betti values of 3.47, 3.08, and 1.64. By contrast, the Euclidean Betti values, 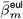, are only 0.42, 0.09, and 0.01. Section 5.2 presents a comprehensive Betti analysis of all activity data. Figure (D-F) analyzes Session 8, Scan 5, from the MICrONS dataset (7).

We quantify how closely neuronal input- and output-similarity matrices, (**C**^in^/**C**^out^), align with functional-similarity matrix (**C**^act^). We compute **C**^in^ and **C**^out^ from the weighted connectome by counting the total number of synapses between each pair of neurons. To capture the network’s global relational structure, we characterize pairwise strength and similarity using these higher-order connectivity measures rather than the direct Hebbian weight matrix (**W**), thereby avoiding additional weighting by synaptic counts or strengths (37–40). For **C**^out^, we include only neurons whole dendrites are fully reconstructed; for **C**^in^, we include neurons whose axons are either extended or clean (7). Functional similarity, **C**^act^, is computed from the full neuronal response across each scan, comprising 30, 000 data points and yielding a median pairwise similarity of 1.18% (Figure 1C). Formally, the connectivity matrices are defined as 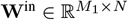 and 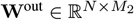., where *N* denotes the number of targeted neurons (or features), and *M*_1_ and *M*_2_ represent the number of structurally confident neurons for axons and dendrites, respectively, where *M*_1_, *M*_2_ ≫ *N* . The connection similarity matrix is then computed as:

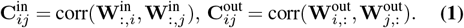

∀ 1 ≤ *i, j* ≤ *N*, **C**^in^, **C**^act^ ∈ ℝ^*N*×*N*^ . We then quantified how each neuron’s connectivity profile relates to its activity profile by correlating the corresponding rows in **C**^act^ with those of **C**^in^*/***C**^out^:

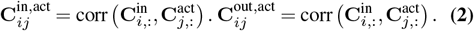

The distribution of connectivity-activity similarities for individual neurons is significantly higher than the shuffled control (Figure 1D1), which revealed a significant dependence of functional similarity on connection similarity. Notably, input similarity is generally a better predictor of functional similarity than output similarity (Figure S2). The neuronal identities and spatial organization of the target neurons are detailed in Table S2, Table S3, and Figure S1. In particular, more than 3*/*4 (382) of the neurons are from V1; the remainder are from higher visual area that are spatially distant. This finding is further supported by centered kernel alignment (Section 4.3 and Figure 1D2) and by analyzing the principal angles between subspaces in the functional and structural affinity spaces (Section 4.4 and Figure S3) (41–43).

### 2.2. Hyperbolic Geometry for Both Connectome and Activity Representation Spaces

Next, we analyze the underlying geometry of the representation space formed by neuronal connectivity and activity using topological data analysis. Specifically, we used Betti curves, which quantify cycles (i.e., closed loops in a topological space) and cliques (i.e., fully connected subgraphs in a simplicial complex) in graphs generated by varying the threshold that defines significant connections between neurons, to identify the underlying geometry of our data (Figure S7) (44). Similar topological methods based on persistent homology has been widely applied to study the manifold structures of population neural or perceptual responses in systems such as olfaction, the hippocampus, head direction, and grid cells, consistently revealing that the observed geometry is often hyperbolic rather than Euclidean (45–49). To estimate the intrinsic dimensionality of the experimental graph, we compare Betti curve distributions from random geometric models of varying dimensionality and select the model that best matches the experimental curve. Specifically, we sample out point clouds in hyperbolic spaces, define similarity as inverse geodesic distance, compute their Betti curves, and quantify agreement with the experimental curves via mean-squared error (MSE) and integrated Betti values. Because Betti curves are invariant under monotone reparameterization, this procedure assesses geometric alignment rather than scale (46, 49). In our data, a low-dimensional hyperbolic model, ℍ^2^, provides the closest match. This analysis indicates that both connectivity and neuronal dynamics are better described by low-dimensional hyperbolic space than with Euclidian space (Figure 1E). The advantages of hyperbolic geometry become more pronounced as dataset size increases, reflecting its higher embedding capacity and efficient accommodation of additional nodes (Figure S16). This result is robust across different connectivity patterns of **W** and across multiple neuronal similarity measures (Section 5.1 and Section 5.3). Additional methodological details are provided in Section 4.2. Finally, applying hyperbolic t-SNE to embed proofread neurons in ℍ^2^ reveals clear cell-type separation, further supporting hyperbolic organization in these circuits (Figure S5) (50). These findings point to a shared geometric substrate linking structural and functional spaces. Notably, low-dimensional hyperbolic embeddings capture the data’s underlying structure with fidelity comparable to that of much higher-dimensional Euclidean embeddings, providing a robust and scalable framework for analyzing and interpreting large-scale neural circuits (Figure S20) (45).

### 2.3. Activity Trace Reconstruction through Linear Model

Next, we tested whether the activity of individual neurons could be directly predicted based on the activity of others and their anatomical connectivity. We implemented a simple statistical linear model, previously shown to be effective in fMRI analysis (51). Given an activity trace **A** ∈ ℝ^*N*×*T*^ with *N* targeted neurons and *T* time points, neuronal dynamics are reconstructed using:

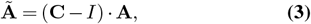

where **C** ∈ ℝ^*N*×*N*^ represents pairwise affinities (i.e. weights of dynamical similarity), and **Ã** ∈ ℝ^*N*×*T*^ denotes the reconstructed activity. We define ***α*** = **C** − **I**, so that ***α***_*ij*_ represents the weight assigned to neuron *i*’s activity trace when predicting the activity of neuron *j* (Figure 2). This model predicts each neuron’s activity by leveraging the simultaneous activity of all other neurons in a leave-one-out manner. To evaluate model performance, we compute the reconstruction accuracy *acc* as the average Pearson correlation between corresponding rows in **Ã** and **A**, enabling neuron-wise comparison. When **C** = **C**^act^, *acc*_act_ reaches 14.1%, averaged across 12 scans. This serves as an upper-bound of all the other linear models, as the model directly uses information from observed activity. To predict activity from structure without accessing the activity of the targeted neuron, **C** can be replaced with either the in-degree (**C**^in^) or out-degree (**C**^out^) correlation matrices derived from the connectome (Section 2.1), allowing structural similarity to serve as the predictor. Alternatively, instead of using the full connectomic, samples can be projected into a low-dimensional space using non-metric multidimensional scaling (MDS) to either hyperbolic or Euclidean space (50, 52). Notably, activity correlation varies substantially across scans due to changes in the mouse’s behavioral state during visual stimulus presentation. This results in significant variability in absolute reconstruction accuracy, largely driven by a small subset of neurons (36). To mitigate this effect, we focus on relative performance, quantified as the explanation ratio *acc/acc*_act_, which measures the effectiveness of alternative affinity measures relative to **C**^act^ (20).

**Fig. 2.**
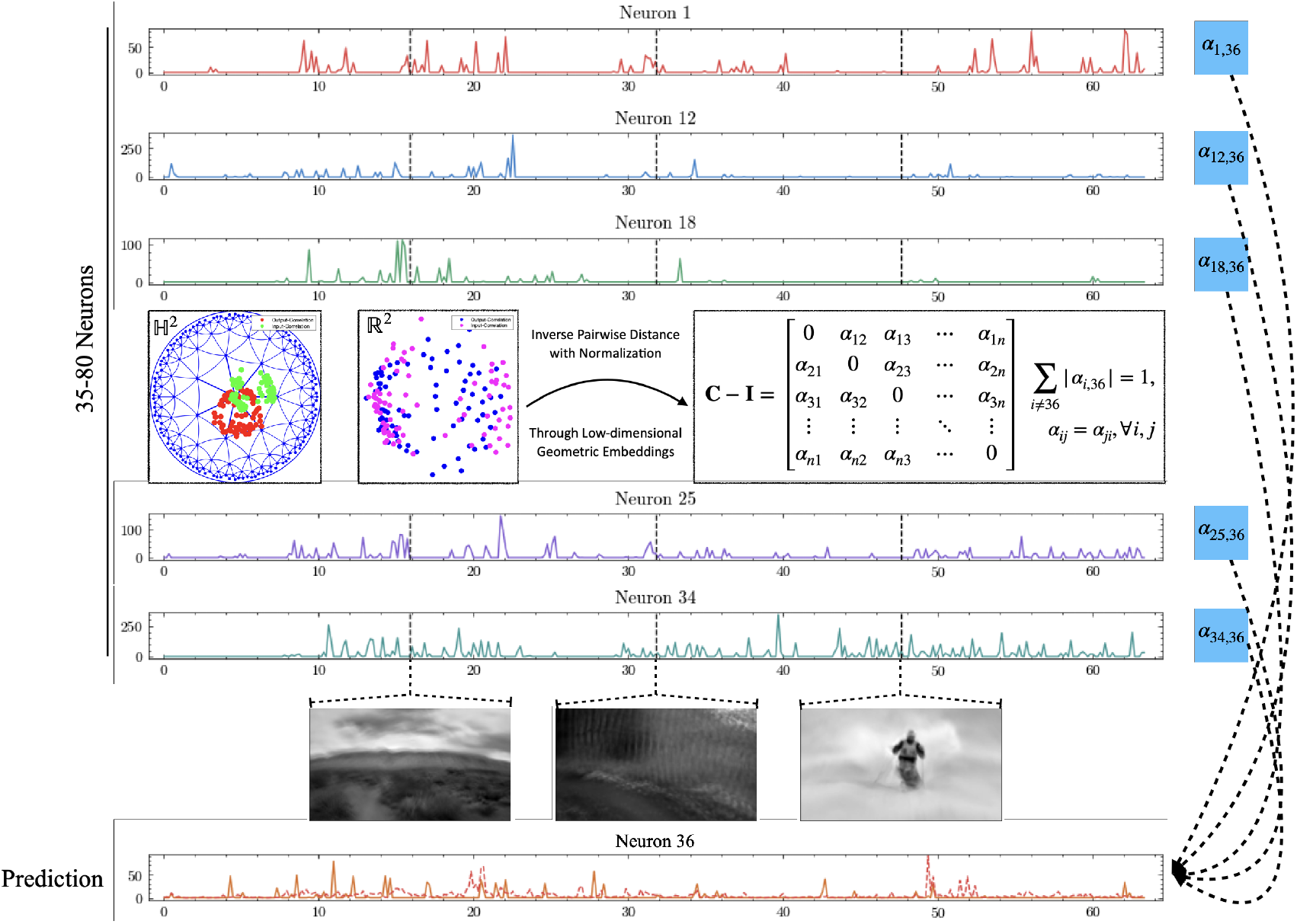
Schematics of Linear Model: The model utilizes deconvolved calcium traces from coregistered imaged neurons recorded during serial video trails to predict the activity of a held-out neuron (7, 36). The prediction is obtained by computing a weighted sum of activities from other neurons, with weights represented by the symmetric matrix **C**. We present 1% of the total traces from Session 8, Scan 5 of the MICrONS dataset. Solid line: groundtruth; dashed line: prediction. Frame rate: 6.3*/s. x*-axis in seconds. Inset: MDS projections of **C**^in^ and **C**^out^ to low-dimensional Hyperbolic and Euclidean spaces.

Using the input connectome correlation, we observe an explanation ratio of 56.3%, whereas the output connectome yields 35.6% (Figure 3A). The observation that **C**^in^ yields better accuracy than **C**^out^ is consistent with the discussion in Section 2.1. Furthermore, having more neurons included in the scan (larger *N* ) improves reconstruction accuracy when using connectome or activity similarity, suggesting that further experimental refinement may improve performance (Figure S21). However, the explanation ratio remains stable and uncorrelated to *N*, highlighting the robustness of the observation (Figure S22). Moreover, we evaluate the performance of direct synaptic connectivity – another common approach to quantifying ***α*** – within the linear model and find that it performs significantly worse than input similarity, although comparably to output similarity, regardless of the connectivity modality (Figure S33) (20, 53).

**Fig. 3.**
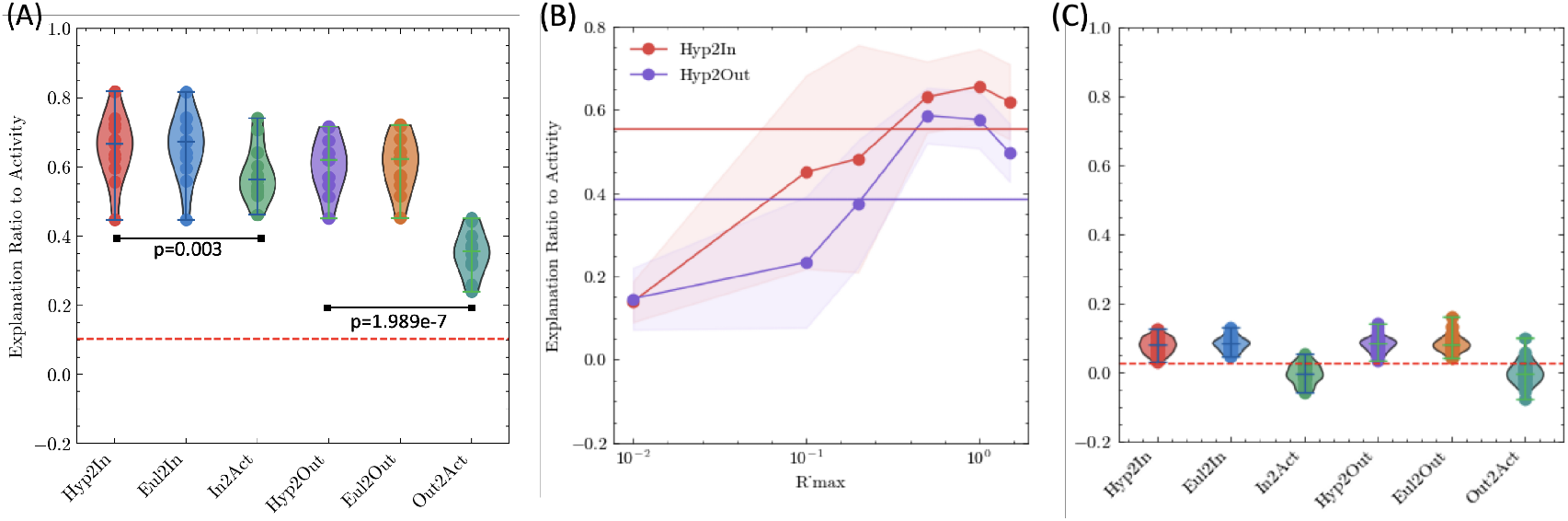
Using Low-Dimensional Geometric Embeddings of Structural Relationship Improves Reconstruction Quality of Activity Traces: (A) Explanation ratio (normalized by scan-specific *a*_act_) achieved by **C**^in^, **C**^out^, and their respective 2D Euclidean and hyperbolic embeddings (*R*_max_ = 1). The dashed line (15.1%) indicates the average of the median pairwise neuronal activity correlations across 12 scans, normalized by *a*_act_. The *p*-values, calculated from paired (dependent) t-test, comparing the performance of **C**^in^ and **C**^out^ with their corresponding hyperbolic embeddings are shown. Each scattered point corresponds to a separate scan and represents the median performance of scan-specific targeted neurons; (B) Sensitivity analysis of reconstruction accuracy as a function of embedding space capacity, varied through *R*_max_. Horizontal lines represent median accuracy for **C**^in^ and **C**^out^; (C): Similar analysis as in (A), but on an untrained continuous-time recurrent neural network, where no signal is represented through the connectome and minimal information is derived from its embeddings.

Surprisingly, utilizing the embedding results from either geometry improves reconstruction accuracy compared to using the raw connectome correlation (e.g. for hyperbolic embedding, 66.6% for input and 62.0% for output), despite a substantial reduction in the dimensionality of the connectome information (Figure 3A). The stability range of *R*_max_ is relatively broad (Figure 3B). The explanation ratios for hyperbolic and Euclidean embeddings are statistically similar across scans, likely due to the limited number of coregistered neurons, which reduces structural complexity and introduces a selective bias, resulting in a nonuniform or bimodal distribution caused by nonhomogeneous neuronal identity (Section 5.1 and Figure S8). Only neurons that are both reconstructed and coregistered are included in the structure-function analysis, whereas all reconstructed neurons are used for the hyperbolic geometry analysis (Section 4.1). Thus far, we exclusively focused on the 2D embedding. When we increased the embedding dimension and surprisingly observed that, despite significant differences in compression quality, the overall reconstruction remained well aligned, underscoring the model’s robustness (Figure S32).

Our results remain generalizable and stable when **C**^in^ and **C**^out^ are computed using either the binary connection pattern or postsynaptic density (PSD) volume (Figure S23), or when considering connections exclusively within subpopulations of specific cell types (Figure S24) (54, 55). Compared to the baseline experiment, using the binary connectome yields comparable accuracy in activity reconstruction (56.2% for input and 36.0% for output), whereas incorporating PSD information significantly reduces performance (52.0% and 28.0%, respectively). Furthermore, even partially accurate structural information – specifically, incorporating connections between unproofread E/I neurons in the computation of **C**^in^*/***C**^out^ – remains beneficial for activity prediction. The most notable improvement is observed for **C**^in^ (64.6%), with a marginal improvement for **C**^out^ (37.8%) (Figure S25). Embedding-based results remain relatively consistent across different connectivity representations. Additionally, restricting connections from or to certain subpopulations of neurons (e.g. E/I) significantly reduces the dimensionality of **W** and limits the performance (36.4% for excitatory input and 48.8% for inhibitory input), whereas the embedding results are less sensitive to this redution (63.8% for excitatory input and 66.5% for inhibitory input) (Table S1, Figure S24) (39, 56).

To test whether the structure–function coupling uncovered by our embeddings generalizes across distinct sensory contexts and even at the raw-signal level, we next dissect prediction performance within individual stimulus periods and for unprocessed fluorescence traces. Although our initial analysis considered neuronal activity across the entire scan, multiple stimulus types were presented in discrete periods (Section 4.1).

To accommodate this structure, we compute **C**^act^, reconstruct activity traces separately within each period, and report the averaged performance. As expected, explanation ratios decrease decline because **C**^act^ now captures more period-specific structure, yet embeddings still confer a clear advantage (see *p*-values in Figure S26). For completeness, we also evaluate the model directly on the raw fluorescence data, which exhibit substantial population-level pairwise correlations (median of 4.90% and mean of 7.00%) (7). The benefit of input-based embeddings become less significant, whereas the advantage of output-based embeddings is retained (see *p*-values in Figure S27 and Table S5).

To test how information-efficient and robust our structure–function mapping truly is, we first ask how prediction accuracy degrades when only a fraction of the available neurons or connections is retained. We explore downsampling by selecting neurons either based on the top-K entries or at random. Specifically, after computing ***α***, we filter (zero out) its entries column by column, either by magnitude or randomly, and then predict the leave-one-out neuron using only a subset of the *N* neurons (Figure S28). Notably, **C**^act^ preserves its accuracy when using only the top-20%, whereas the performance of structurally derived representations deteriorates significantly, particularly for embedded results. For example, in the baseline experiment (Figure S28A), selecting the top 20% entries from **C**^act^ preserves 93.0% of its performance, while only 70.8% and 85.1% are retained for **C**^in,hyp^ and **C**^in^ respectively. The degradation of **C**^in,hyp^ arises because subsampling reduces data quantity, and embedding further compresses dimensionality, together incurring greater information loss compared to the higher resolution and structural density of the original array (**C**^in^). The performance advantage of **C**^in,hyp^ over **C**^in^ becomes less pronounced with smaller values of *K* (*p* = 0.003 for 100% (Figure 3A), 0.051 for 50%, and 0.905 for 20%). However, the significance level is preserved under random selection (*p* = 0.001 for random 50% and 0.002 for random 20%). The observation remains qualitatively consistent across different models (Figure S28). Furthermore, we evaluate the robustness of the embedding results when the connectome (**W**) is corrupted by randomly permuting synaptic connections (Section 5.6). Output correlation exhibits a 19.3% performance degradation when 40% synapses are perturbed, compared to the baseline experiment, whereas the hyperbolic embedding shows only a 1.7% decrease (Figure S29). This highlights the model’s robustness in the presence of unobservable experimental errors.

To further refine our analysis, we restrict the dataset to neurons that are both functionally coregistered and fully proofread (*N* = 338 in total). This filtering enhances reliability by leveraging output similarity in prediction and generally improves relative performance compared to using the unfiltered dataset, partly because the performance of **C**^act^ also declines for small *N* (Figure S21 and Figure S30). To maintain consistency with the previous geometric analysis (Section 5.1), we perform MDS on all proofread excitatory neurons (*N* = 712) to obtain a universal embedding that preserves relevant interactions across a large subset of neurons. We then extract the corresponding information for the scan-specific coregistered neurons for reconstruction purposes.

We also construct a null model using the classical continuous-time recurrent neural network (RNN) (57). Without any training, we inject random inputs and compute the correlation of hidden activities in the recurrent layer with 60 neurons. To ensure a fair comparison with the MICrONS dataset, we finetune the inputs so that the dimensionality of the resulted activity space, quantified through normalized participation ratio, matches that of the experimental data ( ≈ 0.30) (Figure S31) (58). To mimic the typical non-negative distribution of deconvolved activity traces (Figure 2), we use ReLU as the activation function, which naturally constrains values in **C**^act^. Notably, projecting random matrices into a low-dimensional space presents empirical challenges, often causing pairwise distances to approach the space limit (Figure S20), leading to a high stress value. Figure 3C illustrates the correlation of the reconstructed results, where **C**^act^ achieves promising performance with ≈ 74% accuracy. In contrast, both **C**^in^ and **C**^out^ follow an approximately Gaussian distribution centered at 0, effectively nullifying the alignment effect through total summation. The weak signal observed in the embedded results can be attributed to the averaged signal from population activity. This comparative study of the random network highlights that biological systems provide a well-structured scaffold with meaningful structure-function alignment, which can be leveraged to infer and adapt information across different levels.

## 3: Discussion

In this work, we investigate the relationship between synaptic connectivity and the functional responses of excitatory neurons in the mouse visual cortex, leveraging dense reconstruction data from the MICrONS dataset (7, 36, 59). Our analysis reveals a clear alignment between pairwise structural and functional affinities: neurons that receive similar inputs or outputs tend to exhibit similar functional behaviors, and vice versa. Furthermore, by identifying matching topological signatures, we demonstrate that both structural and functional representation spaces can be embedded in low-dimensional hyperbolic spaces with varying radii. Using non-metric MDS method, we construct a correlation matrix based on the spatial proximity of points within these geometric embeddings. Notably, using these results, we achieve not only comparable but significantly improved reconstruction accuracy of activity traces via a simple linear model, despite operating in a significantly lower-dimensional space than the full connectivity matrix. This observation suggests that many anatomical details may not be critical for understanding coarse structure-function mapping and that their interactions can be effectively captured in a reduced parameter space.

It is well established that excitatory neurons with similar responses properties are more likely to be synaptically connected in the mouse primary visual cortex (39, 40, 60–65). Consistent with this, we also observed like-to-like connectivity in our analysis (Figure S4). This phenomenon is proposed to arise in networks where enhanced connectivity strengthens functional covariance and where learning promotes similar responses to similar stimuli (36, 66). Many studies have demonstrated a strong relationship between direct connectivity among targeted neurons (**W** ∈ ℝ^*N*×*N*^, as opposed to **C**^in^ or **C**^out^) and functional interaction, as shown in human mesoscale projectomes, macaque axonal tract tracing study and mesoscale study in Drosophila (20, 53, 67, 68). More similar to (69), we see that structural similarity is a better predictor of activity than direct synaptic connections, since the consideration of non-local interactions increases the fraction of all potential inputs and drivers, rather than restricting to each neuron’s immediate casual inputs (Figure S33). This broader scope could enable the identification of additional structure-function relationships in meso- and macro-scale brain networks (23). Additionally, using direct connectivity significantly limits the amount of data available for reconstruction and constrains performance, as the overall MICrONS dataset exhibits a sparsity of only 0.12%, and the pairwise connections among our 497 targeted neurons have a sparsity of 1.75% (7). In contrast, the correlation measure, acting as a smooth filter, substantially broadens the scope of usable information and produces bounded and symmetric results, making it more suitable for embedding-based analyses. It also enables detailed characterization of preferential selectivity for both inputs and outputs separately (36, 38). Nevertheless, our approach does not address higher-order functional motifs or more complex organizational principles beyond pairwise rules, nor does it account for global topological features such as modularity, efficiency, or short path lengths (36, 69, 70).

Our findings demonstrate that weighted connectivity patterns yield activity prediction accuracy comparable to that achieved using binary connectivity, consistent with prior studies suggesting that the topological structure, defined by the adjacency matrix, is sufficient for structure-function prediction (36, 69).

Also, the use of unweighted graphs to model excitable dynamics also offers computational advantages by simplifying network representation and analysis while maintaining high performance (71). Additionally, our results suggest that total synaptic strength, measured as the sum of PSD sizes, may not serve as a reliable indicator of such alignment (Figure S23). For improved interpretability, we employ a simple linear model to reconstruct activity traces. However, more complex neural-network-based models that incorporate nonlinearity and leverage embedding information, or that simply make ***α*** parameters trainable, such as a single-layer perceptron, may yield improved results (31, 32, 57). Also, considering the design of neurophysiological experiments in which a head-fixed mouse freely walks on a cylindrical treadmill while viewing visual stimuli, the effectiveness of our simple model occurs may result from the passive nature of stimuli presentation, as the mouse is not engaged in any demanding task such as decision making based on contextual information or cues (Figure S31) (7, 20, 72). Evaluating the model’s effectiveness and robustness under various experimental settings represents an interesting direction for future research. Also, oracle scores for functionally coregistered neurons are provided in the MICrONS dataset. This reliability metric is derived from neural responses to repeated natural video stimuli and is computed using a jackknife correlation approach (Figure S34) (7). A low oracle score indicates that a neuron’s activity is inconsistent or stochastic in response to the same stimulus. However, this metric appears uncorrelated with the neuron-wise predictability defined by our model (Figure S35); notably, the activity of a neuron with a low oracle score can still be partiality explained by other neurons. We leave the integration of this feature into the model – for example, using it to differentiate or exclude neurons – to future work. Additionally, we focus on activity reconstruction, noting that subtle modes of activity may play a critical role in behavior (73). Our work exclusively examines the recurrent circuit components reconstructed in the MICrONS dataset and does not make any assumptions or provide formulations regarding how stimuli are presented or transmitted through the sensory cortex (e.g. retinal ganglion cell (RGC) and lateral geniculate nucleus (LGN)) (36, 59, 74).

In this study, we demonstrate that both the structural connectome and the corresponding functional activity patterns of neurons can be embedded within the same low-dimensional hyperbolic space. The exponential volume growth and low distortion characteristic of hyperbolic geometry enable compact, efficient representations that are especially well-suited to the hierarchical, tree-like organization often found in biological networks (75, 76). This structural efficiency reduces storage and computational costs while supporting scalable, interpretable embeddings of complex neuronal data. Moreover, hyperbolic geometry underpins robust communication protocols that rely only on local network information, preserving efficiency even as connectivity dynamically changes (77, 78).

We employ an RNN with randomly initialized weight as a null model, showing that the organized structure captured by **C**^in^ or **C**^out^ is a necessary prerequisite for an embedding to achieve a high explanation ratio. Because the reconstructed excitatory neurons exhibit sparse all-to-all connectivity, we cannot readily incorporate the experimental connectome into simulations to establish sufficiency. In addition, for greater fidelity we compute **C** using rectangular matrices (Section 2.1), whereas the standard RNN formulation assumes a square weight matrix. Obtaining more densely reconstructed experimental data will therefore be crucial for fully addressing this question.

The effectiveness of using MDS to set weights in the linear model may be attributed to at least three factors. First, anatomical similarity may create a spatial embedding for each neuron, where similar neuron types are positioned closer together and exhibit similar activation patterns. This arises from the presence of multiple neuronal types, which are not explicitly differentiated in the recordings (79–81). Second, the spatial location of neurons imposes constraints on connection formation, which MDS then extracts from the connectome (8, 82–84). While the mouse cortex lacks organization in terms of orientation columns, it exhibits strong retinotopic organization (85, 86). These mechanisms are not independent, as spatial distance plays a key role in refining neuronal subtypes and predicting variations in synaptic connectivity patterns (38, 87). Third, neuronal responsiveness and activity patterns may be strongly influenced by the spatial and temporal organization of synaptic inputs, which, in conjunction with Hebbian plasticity, shape synchronous dynamics (39, 60, 88). MDS could effectively capture these dynamics. However, empirical measurements of the dimensionality of MICrONS activity space suggest that neural dynamics are not strictly low-dimensional (Figure S31), limiting the influence of this factor. A precise attribution of MDS effectiveness to these or other factors is beyond the scope of this study.

In conclusion, this study empirically quantifies the structure-function relationship using synaptic-scale connectivity correlations and demonstrates that low-dimensional embeddings of these correlations are more closely associated with neuronal dynamics. Specifically, detailed synaptic information from pairwise neuronal connections can be effectively represented as interactions in two- or three-dimensional spaces with varying geometry, enabled by improved reconstruction efficacy. Accordingly, we emphasize the concept of connectome embedding and its significance for understanding circuit-level mechanisms with broader implications. First, calculating input similarity using only excitatory neurons results in a 20.1% performance decrease, whereas the embedding approach incurs only a 3.11% reduction (Figure 3 and Figure S24). This robustness suggests that connectomic data remains highly informative even without a fully dense, though still unbiased, reconstruction. Second, the comparable performance of the embedded output highlights the potential presence of plasticity rules that generate network configurations amenable to low-dimensional embedding. Third, the effectiveness of connectome embedding indicates resolving every individual synapse may not be necessary to understand structure-activity relations. Alternatively, for example, one could construct a parsimonious, cell-type-based model capable of predicting neuronal connectivity, thereby significantly reducing computational costs in simulations.

## ACKNOWLEDGEMENTS

We thank Kyle Aitken, Stefan Berteau, Yusi Chen, Forrest Collman, Nuno da Costa, Bethanny Danskin, Łukasz Kuśmierz, Zhixin Lu, Dana Mastrovito, Eric Shea-Brown, Uygar Sümbül, and Praveen Venkatesh for discussions. We also wish to thank the Allen Institute for Brain Science founder, Paul G. Allen, for his vision, encouragement, and support.

## 4: Methods

### 4.1. MICrONS Dataset and Experiments

In this paper, all experiments and analysis are conducted using the MICrONS minnie65 V943 dataset, which contains 205,277,810 synapses and 90,230 neurons (7). The overall sparsity of pairwise connectivity is 0.127%. Among the recorded cells, 63,758 are excitatory neurons, 7,849 are inhibitory neurons, and 18,693 are non-neuronal cells, primarily glial cells. Regionally, 57,781 cells are located in V1, 21,184 in RL, and 10,480 in AL. Proofreading plays a critical role in improving the quality of automated EM segmentation by resolving incorrect associations between presynaptic and postsynaptic partners. This process primarily addresses two types of reconstruction errors: false merges and false splits. False merges occur when segmented objects are erroneously grouped (e.g., an axon or dendrite incorrectly assigned to a soma), while false splits arise when objects are mistakenly separated (e.g., an axon or dendrite omitted from its corresponding soma). To correct false merges, the reconstruction is *cleaned* by removing incorrectly associated segments, whereas false splits are addressed by *extending* the reconstruction to restore missing segments. A detailed protocol for morphological proofreading is provided in (7, 36).

For axonal proofreading, 321 neurons are classified as extended, 714 as clean, and the remaining neurons are unproofread (89). Similarly, for dendritic proofreading, 44,163 neurons have been completely merged and properly cleaned – either manually or using the NEURD algorithm – resulting in fully reconstructed dendrites, while the remaining neurons are unproofread (89, 90). To increase the number of axonally trustworthy neurons, we relax the selection criterion to include neurons labeled as either extended or clean. In contrast, reconstructed dendrites are generally complete and require minimal extension, resulting in a greater number of neurons classified as dendritically trustworthy. However, only 845 neurons (fewer than 1% of the total) meet the structural confidence criteria for both axonal and dendritic aspects. In analyzing the geometric representation of the connectome (Section 5.1), we will consider different subsets, including all excitatory neurons (*N* = 716) and excitatory neurons in V1 (*N* = 675). For the analyses in Section 2.1 and the baseline experiment in Section 2.3, **C**^out^ is computed using connections exclusively *to* E/I neurons with fully reconstructed dendrites, whereas **C**^in^ is computed using connections exclusively *from* E/I neurons with extended or clean axons. Table S1 details the dimensionality of **W**^in^ and **W**^out^ under different experimental settings, and Tables S2 and S3 detail the cell identities. Neurons that are isolated in the connectivity are excluded, as their inclusion would result in entire rows and/or columns of NaN values in **C** (corresponding to 0 values in **W**). Additional comparative studies, including those involving different connectome types **W** or the inclusion of synapses *to* or *from* unproofread neurons and glial cells, are discussed in Section 2.3 (Table S1, “Noise” row).

In this study, we align the anatomical and functional properties of excitatory neurons that are both functionally coregistered (i.e. located within the volume of dense reconstruction, also see Figure S9) and structurally proofread. These selection criteria are implemented using CAVE (35). While restricting analysis to neurons that meet both criteria enhances data reliability, it significantly reduces the number of available neurons. Requiring proofreading of both the axon and dendrite would shrink the dataset even further, so we include coregisted neurons if at least one of these compartments has been proofread (**Prf**). Table S4 provides a detailed comparison of selection criterion (36). We use the number of neurons listed under “Number of **Coreg** + **Prf** Neurons” in the linear model presented in Section 2.3. Across all scans, only 497 **Coreg** + **Prf** neurons (excluding duplicate appearances) meet these criteria. During the recordings, three dynamic stimulus types, natural movies, global directional parametric stimuli (“Monet”), and local directional parametric stimuli (“Trippy”), were presented in blocks interleaved with 8–12 off-periods (7, 36).

### 4.2. Clique Topology Method for Uncovering Hidden Geometries in Connectomic and Activity Structures

It has been extensively discussed in previous literature that the clique topology method could be used to detect the geometric organization of the correlation or connectivity matrix, including purely random or embedded in Euclidean or hyperbolic space (45, 46, 49). In this analysis, we extend these discussions by using the input as a symmetric matrix **C**, which quantifies the pairwise correlation between neurons *i* and *j*. The algorithm applies a step function to the matrix at various thresholds, retaining only the entries that exceed the specified value and generating corresponding topological graphs. These graphs are characterized by the number of cycles (holes) in different dimensions (1, 2, or higher). These counts, known as Betti numbers 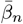, are generally small at low thresholds (fully connected networks) or high thresholds (sparse networks where most nodes are not interconnected) (91). Plotting them as a function over the entire density range *ρ* of links (from the highest threshold to the lowest threshold) produces Betti curves *β*_*n*_(*ρ*), 0 ≤ *ρ* ≤ 1. Clearly:

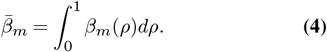

Typically, the first three Betti curves are sufficient to reveal the underlying structure, and our analysis adopts this convention. Figure S7 provides an illustrative example of computing Betti curves for a simple graph.

Uniformly sampling points from an *n*-dimensional Euclidean space is straightforward through high-dimensional uniform distribution sampling. Previous literature has also addressed sampling from *n*-dimensional hyperbolic space (46, 49). In the polar coordinate setting, both the curvature *K* and the maximal radius *R*_max_ can alter the geometry of the space. Specifically, changing *K* only rescales the radial coordinates of the nodes without affecting the topological properties of the network. For simplicity, we fix 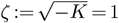 and treat *R*_max_ as a hyperparameter. Each point is sampled with uniformly distributed angles and a radius *r* ∈ [*R*_min_, *R*_max_], following the distribution *ρ*(*r*) ∼ sinh^(*d*−1)*r*^, which is approximately proportional to *e*^(*d*−1)*r*^ when *r* ≫ 1 . In (46), *R*_min_ is set to 0.9*R*_max_ to constrain the sampled points to have similar radii (i.e. similar to torus-like shape in ℝ^3^). Following the convention in (49), we set *R*_min_ = 0, leading to a less constrained distribution. Due to this difference in setting, the Betti curves presented in our work are qualitatively more similar to those in (49) and differ from those in (46) (e.g. decreasing prominence of peaks as *m* increases). Consequentially, unlike the *L*^2^ distance in Euclidean space, the metric in hyperbolic geometry adheres the hyperbolic law of cosines:

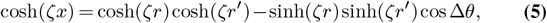

where *x* is the hyperbolic distance, *r, r*^*′*^ the corresponding radii (of the two points) from the origin, and Δ*θ* is the angle in between (49). To incorporate noise into the simulation, we consider the magnitude of Gaussian noise *ϵ*, which is scaled based on the geometric distance 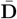 calculated through Equation (5):

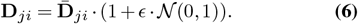

We use the negative distance, **C**_*ji*_ = **C**_*ij*_ = − **D**_*ji*_, to quantify the correlation between neuron *i* and *j*. Any other order-reversing operation, including the reciprocal, could also accomplish this transformation, but we follow the previous literature and use the inverse for its stability in scaling (46, 49). In the main text, we primarily use Pearson’s correlation to calculate **C**. However, any operation that produces bounded and symmetric results can be applied, including Jensen-Shannon divergence, Hellinger distance, Jaccard distance, or Czekanowski-Dice distance for quantifying connectome similarity, and cosine similarity for quantifying activity similarity (83). In Section 5.3, we demonstrate the robustness of the low-dimensional embedding geometry by using cosine similarity as an alternative method to compute **C** for activities. In all simulations, we set *N* to match the size of each distance matrix as the input, with *δ* = 0.1, *ϵ* = 0.0625, *d* = 2 (or 3, see Section 5.4), while fine-tuning the hypodermic radius *R*_max_ to achieve optimal alignment of the top three Betti curves. The optimization objective is to minimize the cumulative absolute pointwise error (Frobenius norm) of Betti curves using experimental and synthetic data. We validate the results by comparing the Betti number, 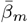. The analysis will display ± 1 standard deviation in all such cases. Through experiments, we observe that both the shape and magnitude of *β*_*m*_(*ρ*) are highly dependent on *ϵ*, which can be interpreted as representing various levels of mixing with random matrices (Equation 6). The parameter values are selected based on prior literature with minor adjustments (46, 49). Benefiting from recent developments, the scope of network size considered in the topological analysis presented in this paper surpasses that of previous works multiple times (Section 4.6).

A key difference between clique topology analysis and hyperbolic MDS (Section 2.3) is that the latter applies dimensionality reduction to empirical data by minimizing the stress function and does not guarantee a uniform distribution of embedded points. The validity of our topological analysis is supported by the relatively uniform distribution of the targeted neurons, as confirmed through Euclidean non-metric MDS, which satisfies the assumptions underlying this analysis (Figure S6) (46, 49). In practice, we initialize the hyperbolic MDS algorithm with the converged Euclidean embedding of the same dimension and observe improvements in stress, highlighting the enhanced embedding accuracy of hyperbolic MDS.

### 4.3. Centered Kernel Alignment

Centered kernel alignment (CKA) gauges how similar two sets of neural activations are by comparing their centered Gram (kernel) matrices (43). Given activation matrices *X* ∈ ℝ^*n*×*p*^ and *Y* ∈ ℝ^*n*×*q*^ for the same *n* features (in this study, the latent dimension of the reduce representation), form linear kernels *K* = *XX*^⊤^, *L* = *Y Y* ^⊤^, and mean-center them with 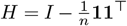 to obtain *K*_*c*_ = *HKH, L*_*c*_ = *HLH*. CKA is the cosine between these two matrices:

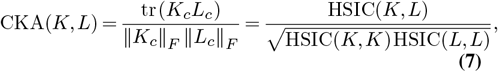

where HSIC is the Hilbert-Schmidt independence criterion. For the common linear CKA variant this reduces to:

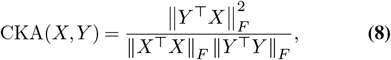

which lies in [0, 1]: values near 1 mean the two representations rank stimuli in almost the same way, whereas values near 0 indicate unrelated structure. Because CKA is invariant to any orthogonal rotation of features and to isotropic rescaling, it avoids spurious differences that plague canonical-corelation-based metrics, and, unlike fully linear-invariant measures, still distinguishes wide layers when *p, q* > *n*.

### 4.4. Principal Angle Analysis

Principal angle analysis quantifies the overlap between two neural-activity subspaces (41, 42). Given column-orthonormal bases *U* ∈ ℝ^*d*×*p*^ and *V* ∈ ℝ^*d*×*q*^, form *U*^⊤^*V* and compute its singular values *σ*_1_ ≤舰 ≤ *σ*_min(*p,q*)_. The principal angles are *θ*_*i*_ = arccos *σ*_*i*_ with 0^°^≤ *θ*_*i*_ ≤ 90^°^. The first angle *θ*_1_ measures the most-aligned pair of directions, whereas subsequent angles describe alignment after removing previously matched components. The average squared cosine, 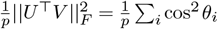 equals the fraction of variance that one subspace projects onto the other. Small angles (near 0^°^) indicate shared coding dimensions, while large angles (near 90^°^) denote independence. Because the computation scales efficiently when *p, q* ≪ *d*, principal angle analysis is widely used to compare population codes across tasks, time points, and brain regions (92).

### 4.5. Linear Model

In the baseline experiment, glial cells are excluded to focus on interactions between excitatory (E) and inhibitory (I) neurons. Synaptic sign is assigned according to Dale’s law, with inhibitory neurons having negative output weights and excitatory neurons having positive output weights. Neuronal identity is determined based on morphological features (7). **C**^in^ and **C**^out^ are computed using the same approach described in Section 2.1. Synaptic information from all functionally coregistered neurons across scans (*N* = 497), including neurons without complete structural confidence, is concatenated and subjected to dimensionality reduction. To avoid overfitting, we bin the activity data, compute **C**^act^ using the concatenated information from odd-numbered bins, and evaluate *acc*_act_ on the remaining data.

While the effectiveness of using correlations derived from the input connectome are intuitive, the ability of the output connectome correlations to predict activity is less immediately clear (18, 93). Previous studies employed a “digital twin” model to predict neural responses to a diverse stimulus set of natural and parametric movies, characterizing the *in silico* functional properties of imaged neurons. These studies revealed an overarching like-to-like principle – neurons with highly correlated responses to natural videos tend to be connected across multiple layers and visual cortical areas – providing insights into the effectiveness of **C**^out^ (36, 59, 94).

### 4.6. Code availability

All analysis code for this study is publicly available on GitHub^1^. We also adapted the code from pervious works on topological data analysis here^2^, hyperbolic MDS^3^, and hyperbolic t-SNE^4^ in this work.

## 5: Supplementary Materials

**Table S1.**
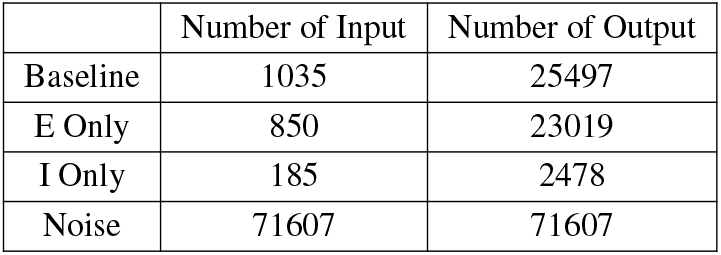
Number of Input and Output Neurons Used in Calculating C^in^ and C^out^: Corresponds to the values of *M*_1_, *M*_2_ . All rows, except for “Noise”, include only neurons that are proofread on the axon side (for input) and the dendrite side (for output) correspondingly.

|  | Number of Input | Number of Output |
| --- | --- | --- |
| Baseline | 1035 | 25497 |
| E Only | 850 | 23019 |
| I Only | 185 | 2478 |
| Noise | 71607 | 71607 |

**Table S2.** Layer Classification of Input, Output, and Target Neurons in the Baseline Experiment: L: Layer; WM: White Matter.

|  | L1 | L2-3 | L4 | L5 | L6 | WM |
| --- | --- | --- | --- | --- | --- | --- |
| Input | 10 | 407 | 355 | 182 | 77 | 4 |
| Output | 402 | 7751 | 7453 | 3828 | 4719 | 1344 |
| Targeted | 1 | 205 | 157 | 92 | 42 | 0 |

**Table S3.** Region Classification of Input, Output, and Target Neurons in the Baseline Experiment: V1: Primary Visual Cortex; AL: Anterolateral Area; RL: Rostrolateral Area.

|  | V1 | RL | AL | Others |
| --- | --- | --- | --- | --- |
| Input | 899 | 88 | 48 | 0 |
| Output | 18895 | 6158 | 413 | 31 |
| Targeted | 382 | 68 | 47 | 0 |

**Fig. S1.**
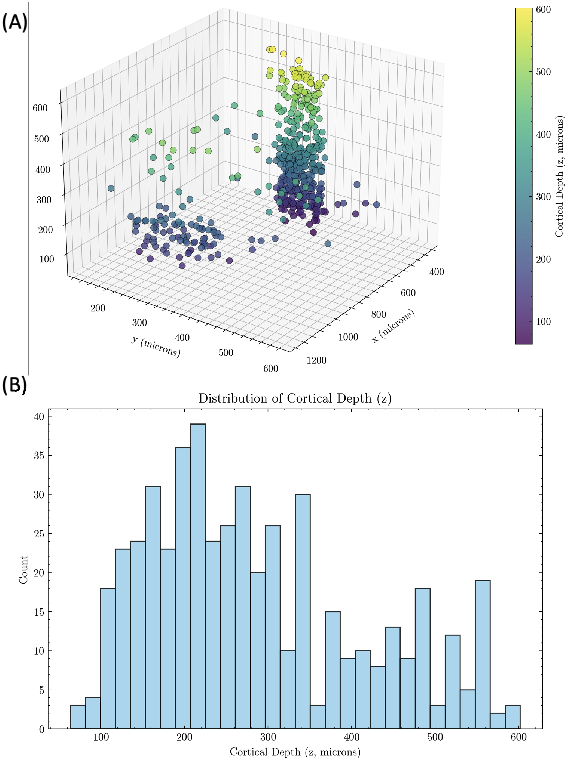
Spatial Organization of Target Neurons: (A) Somatic locations of the 497 targeted neurons, color-coded by cortical depth (*µm*); The *z*-coordinate should be flipped in order to be aligned with the physical orientation; (B): Histogram of cortical depth. Results are consistent with Table S2 and Table S3.

### 5.1. Representation Space of Neuronal Connectome is Low-Dimensional Hyperbolic Embedded

See Figure S8, S9, S10, S11, and S12.

**Fig. S2.**
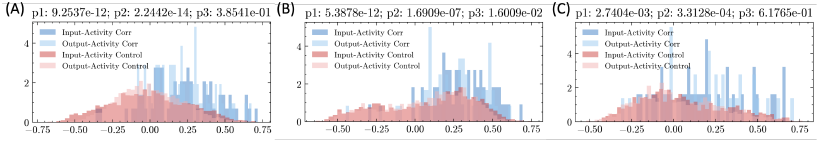
Statistical Comparison of On-Diagonal and Off-Diagonal Elements of “Correlation of Correlation” Matrix: Expanding on Figure 1D1, we present results from additional scans to demonstrate the consistency and robustness of our observations. (A): Session 4, Scan 7; (B) Session 9, Scan 4; (C): Session 5, Scan 3. The *p*_1_ to *p*_3_ values, defined identically to those in Figure 1D1, are provided in the captions.

**Fig. S3.**
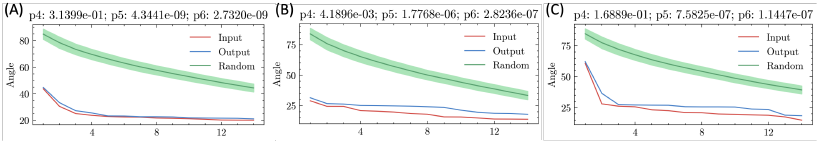
Principal Angle Analysis: Expanding on Figure 1D2, we compute the principal angle between subspaces (flats) constructed by **C**^act^ and **C**^in^ (red), **C**^out^ (blue), and random matrices (green) (41, 42). Statistical significance is indicated by *p*_4_ for the difference between the red and blue curves, *p*_5_ for the red and green curves, and *p*_6_ for the blue and green curves. As discussed in the main text, the input affinity subspace exhibits the strongest alignment with the activity space. (A) Session 4, Scan 7; (B) Session 9, Scan 4; and (C) Session 8, Scan 5 (Figure 1).

**Fig. S4.**
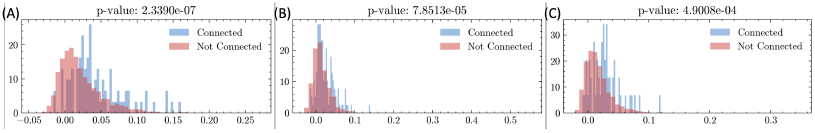
Neurons with Strong Activity Correlation Are More Likely to Form Synaptic Connections: Unlike Figure 1, Figure S2, and Figure S3, which use non-local connectivity measures (**C**^in^ and **C**^out^) to probe the structure-function relationship, here we analyze the direct synaptic connectivity matrix between targeted neurons (**W** ∈ ℝ^*N*×*N*^ ) and confirm the like-to-like principle (36). (A) Session 4, Scan 7; (B) Session 9, Scan 4; and (C) Session 8, Scan 5.

**Fig. S5.**
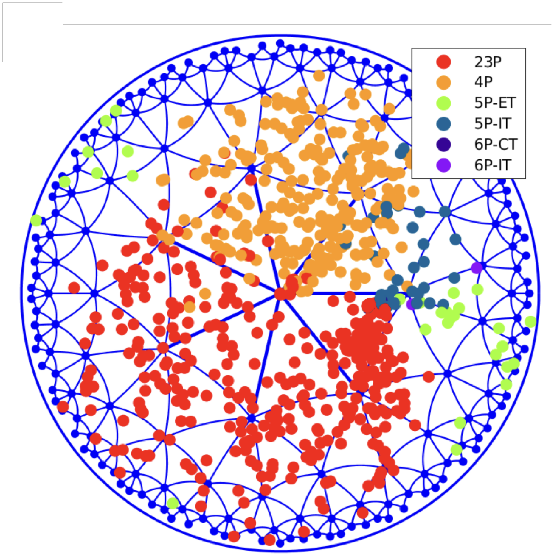
Embedding neurons from Figure 1B based on their output similarty into ℍ^2^ with hyperbolic t-SNE algorithm, clustered by cell type and cortical layer: The resulting projection is displayed on a unit circle (50).

### 5.2. Additional Betti Curve Analysis on Representation Space of Activities Traces for All Scans

See Figure S15 and S16.

### 5.3. Betti Curve Analysis on Representation Space of Activities Traces Using Cosine Similarity

See Figure S17 and S18.

### 5.4. Equivalence Between 2D and 3D Hyperbolic Embeddings

See Figure S19. A similar discussion in Euclidean space is well addressed through the classical Johnson–Lindenstrauss lemma, which establishes the existence of a linear map *f* : ℝ^*N*^ → ℝ^*n*^, *N* > *n*, that nearly preserves the distances between points (95, 96). Here, we shift the focus from pairwise distance preservation to topological feature preservation and observe a perfectly linear relationship in the hyperbolic radius under pointwise transformation from ℍ^3^ to ℍ^2^. Our goal is not to prove the universality of such transformations but rather to highlight the embeddings in ℍ^2^ discussed previously, including Section 5.1, 5.2, 5.3, have direct counterparts in ℍ^3^, as is typical in prior literature (46, 49). The range of 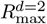 shown in Figure S19 aligns with the discussions in earlier sections.

**Table S4.** Available Neurons under Different Selection Criteria, Categorized by Session and Scan: **Coreg:** functionally coregistered. **Prf:** proofread on at least one side (axon or dendrite).

| Session & Scan | Number of Recorded Neurons | Number of **Coreg** Neurons | Number of **Coreg + Prf** Neurons |
| --- | --- | --- | --- |
| Session 4, Scan 7 | 8395 | 1018 | 82 |
| Session 8, Scan 5 | 10579 | 1123 | 68 |
| Session 6, Scan 4 | 8826 | 1152 | 59 |
| Session 9, Scan 4 | 8415 | 1027 | 56 |
| Session 6, Scan 6 | 8551 | 1052 | 54 |
| Session 5, Scan 6 | 9254 | 997 | 54 |
| Session 5, Scan 7 | 8862 | 1000 | 41 |
| Session 6, Scan 2 | 8756 | 1021 | 39 |
| Session 7, Scan 3 | 9296 | 743 | 38 |
| Session 7, Scan 5 | 9034 | 631 | 39 |
| Session 9, Scan 3 | 8661 | 1170 | 37 |
| Session 5, Scan 3 | 8790 | 426 | 35 |

**Fig. S6.**
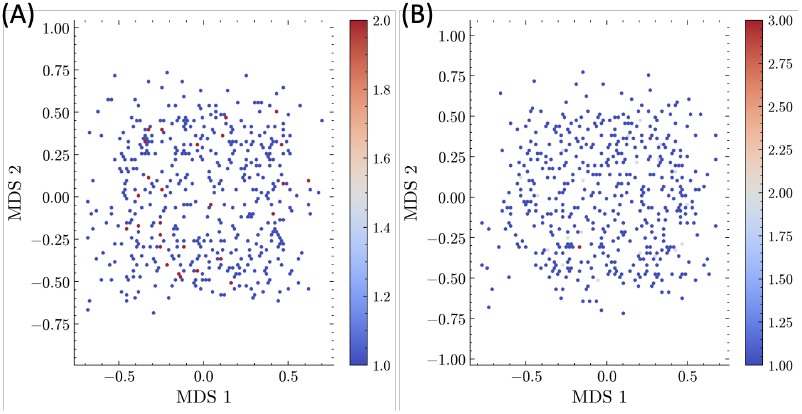
2D Euclidean MDS Embeddings of Input and Output Dissimilarity in the Baseline Experiment: The dissimilarity is calculated through 1 - Pearson correlation. Colored by frequency.

**Fig. S7.**
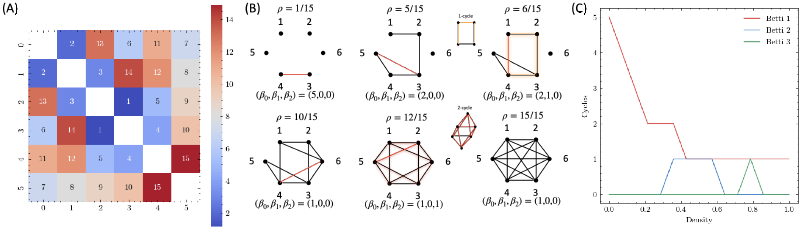
Illustrative example of order-based analysis of symmetric matrics: (A): An arbitrary correlation matrix where each cell’s value is converted to its relative rank; (B): By iteratively adjusting the threshold (filtration density), a sequence of binary adjacency matrics (and graphs) is generated. *β*_0_ counts the number of connected components, while other *β*_*m*_ values count the number of *m*-cycles, such as 1-cycles (squares), 2-cycles (octahedra), and 3-cycles (orthoplex, not shown), which may be created, persist, and disappear at different densities. The colored shadow suggests the presence of a cycle; (C): The overall Betti curves *β*_*m*_(*ρ*) are shown by combing all the discrete instances discussed in (B). This example is reproduced from Dr. Nicole Sanderson’s tutorial at IPAM’s Mathematical Approaches for Connectome Analysis Workshop.

**Fig. S8.**
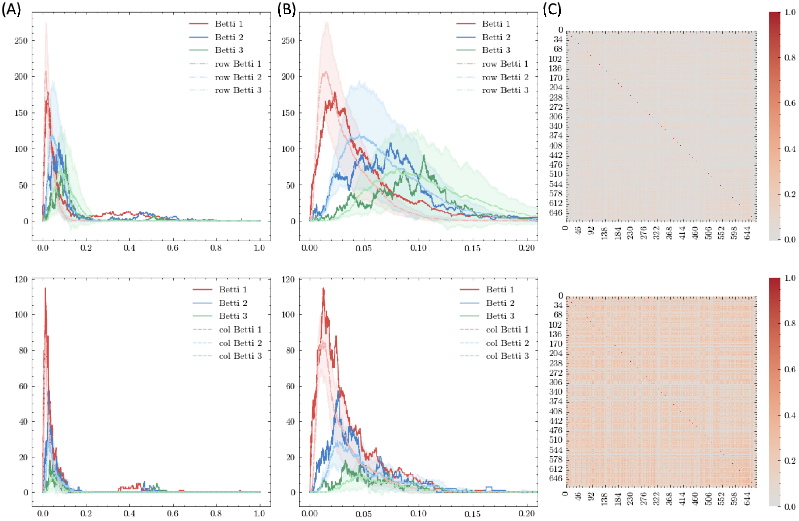
2D Hyperbolic Embedding of Structural Representation of Binary Connectome Among Proofread Excitatory Neurons in V1: To compare with Figure 1E in the main text, square Pearson correlation matrices of the binary connectome are computed for all proofread excitatory neurons in V1 (*N* = 675). As discussed in Section 2.3, structurally incomplete (unchecked) yet informative data is included. **C**^out^ (top row) and **C**^in^ (bottom row) are calculated using the rectangle matrix that includes all recorded excitatory and inhibitory neurons from MICrONS Version 943(7). The reconstructed neurons are distributed across two distinct cylindrical regions, corresponding to the V1 area (94.0%) and the RL/AL area (6.0%). Overall, their structural similarities, both for inputs and outputs, form two clusters, violating the assumption of uniform distribution in Betti analysis (49). Here, we focus specifically on V1 neurons; however, the results remain consistent for RL/AL neurons (Figure S12). A finer classification that groups neurons by cell types produces smoother and more regular Betti curves, thereby achieving better alignment with hyperbolic geometry in terms of topological feature mapping (Figure S13 and S14): (A): Betti curves for groundtruth and synthetic data, sampled from a 2D hyperbolic space (see Section 4.2), averaged across 10 trials due to the high computational memory costs; results are shown with best fits via (pointwise) mean squared error (MSE) comparison to the mean curve (dashed lines) with optimized *R*_max_ of 18 and 14 respectively (grid size = 1); (B): Zoomed-in view of (A) highlighting the low edge density region; (C): Heatmap visualization of each correlation matrix. Using alternative correlation measures instead of Pearson correlation yields consistent results (not shown), as discussed in Section 4.2 and (83).

### 5.5. Reconstruction of Raw Fluorescence Traces

See Figure S27 and Table S5.

**Fig. S9.**
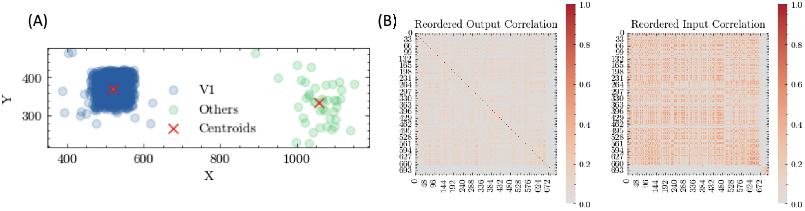
Spatial Distribution of Reconstructed Neurons: (A): Cross-sectional projection on the XY plane (in *µm*) of all reconstructed neurons (*N* = 716), clustered using the K-means algorithm (7); (B): Sorted **C**^out^ and **C**^in^, arranged according to the labels in (A), and the matrices correspond to Figure S8.

**Fig. S10.**
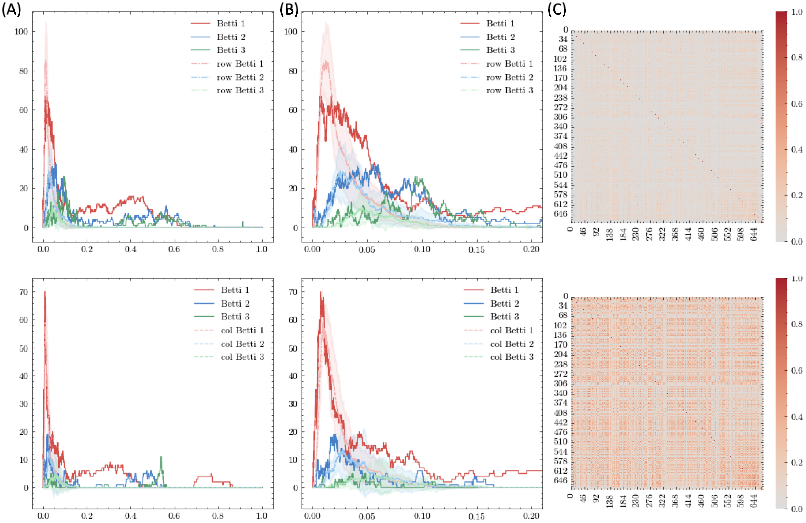
2D Hyperbolic Embedding of the Structural Representation of the Weighted Connectome Among Proofread Excitatory Neurons in V1: Similar to Figure S8, but **C**^out^ and **C**^in^ are computed using synaptic counts between neuronal connections, with optimized *R*_max_ of 14 and 13, respectively. As in Figure S8, the distinct emergence of a second peak in the high edge density region is consistent with observations in (46) and may be attributed to the significant nonuniform distribution of points in ℍ^*n*^.

**Fig. S11.**
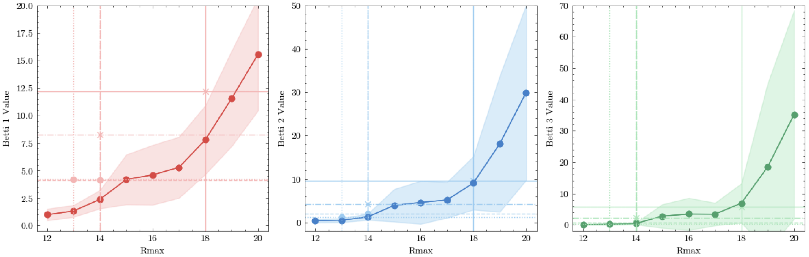
Direct Comparison of Betti Values 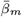 Based On Groundtruth and Synthetic Data: Results derived from Figure S8 and Figure S10; Vertical lines indicate the optimized *R*_max_ values for **C**^out^ and **C**^in^ (4 in total, for binary and weighted connectome respectively); Horizontal lines represent the 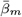 values, *m* = 1, 2, 3, calculated based on the groundtruth. The deviations from the intersection points (“∗” for **C**^out^ and “o” for **C**^in^) to the shadows indicate the effectiveness of approximations. All results are well-aligned except Betti 1 due to the occurrence of the second peaks at the high edge density region. The results further demonstrate the accuracy of the fits.

### 5.6. Using Embedding Results Demonstrates Robustness in Prediction Under Raw Connectome Perturbations

See Figure S29. Here, we perturb the connectivity matrices using the baseline experimental setting but with binary connections, concatenating all targeted neurons across scans (*N* = 497), **W**^in^ and **W**^out^ (Section 2.1). A *k*% perturbation in Figure S29 refers to the random deletion of *k*% of synapses (i.e. false negative entries) and the simultaneous addition of *k*% of “fake” synapses (i.e. false positive entries). This approach preserves the overall connection sparsity (3.87% for **W**^in^ and 0.52% for **W**^out^). The perturbation is subject to the following constraints: 1) It must not result in any isolated targeted neuron, as this would lead to undefined correlation with others, rendering their activity unusable for reconstruction (i.e. undefined ***α*** entries). Omitting neurons could bias the comparison (Figure S21); 2) It must satisfy Dale’s constraint for **W**^in^; since all targeted neurons are excitatory, **W**^out^ only contains 0 and 1 entries. Due to constraint 1) and the limited number of axonally proofread neurons (Table S1), perturbation of **W**^in^ will be biased, therefore Figure S29 presents results only for **W**^out^.

**Fig. S12.**
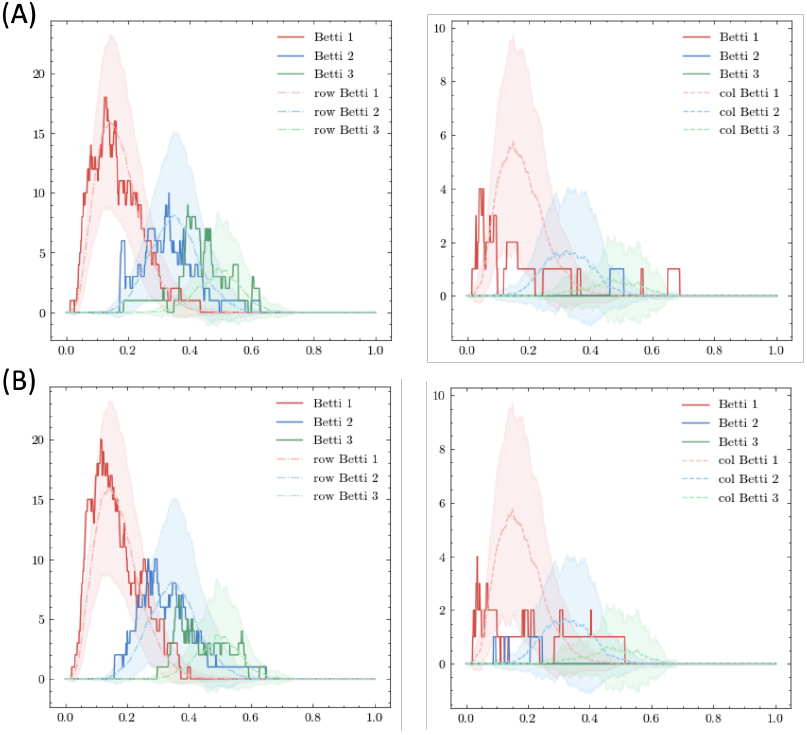
2D Hyperbolic Embedding of the Structural Representation of the Weighted Connectome Among Proofread Excitatory Neurons in RL/AL: Similar to Figure S8 and S10, this analysis focuses on neurons in higher visual areas (RL/AL) (*N* = 41) using (A) binary connectome or (B) weighted connectome to evaluate **C**^out^ (left column) and **C**^in^ (right column). Synthetic data are presented with optimized *R*_max_, averaged across 100 trials. Similar to observations in V1, the topological features of the representation space formed by **C**^in^ are less pronounced and can be approximated using a small *R*_max_, making it more similar to Euclidean space. This may be attributed to neuronal axons traveling much greater distances than dendrites, or alternatively, to an imbalance in reconstruction.

**Fig. S13.**
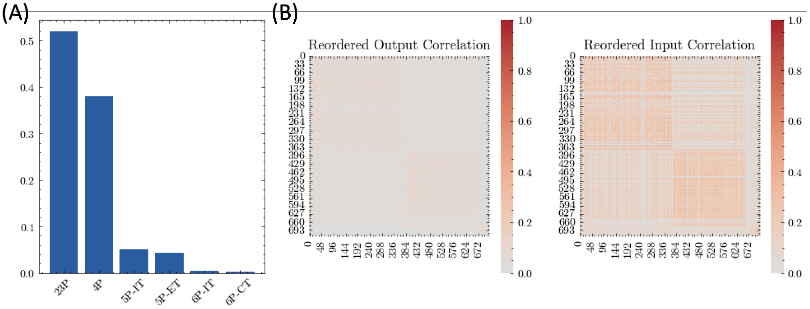
Cell Type of Reconstructed Neurons: (A): Labeled cell types of all reconstructed neurons, where layer 2-3 pyramidal cells (23P, 52.0%) and layer 4 pyramidal cells (4P, 38.0%) are dominant (7); (B): Sorted **C**^out^ and **C**^in^, arranged according to the labels in (A).

**Fig. S14.**
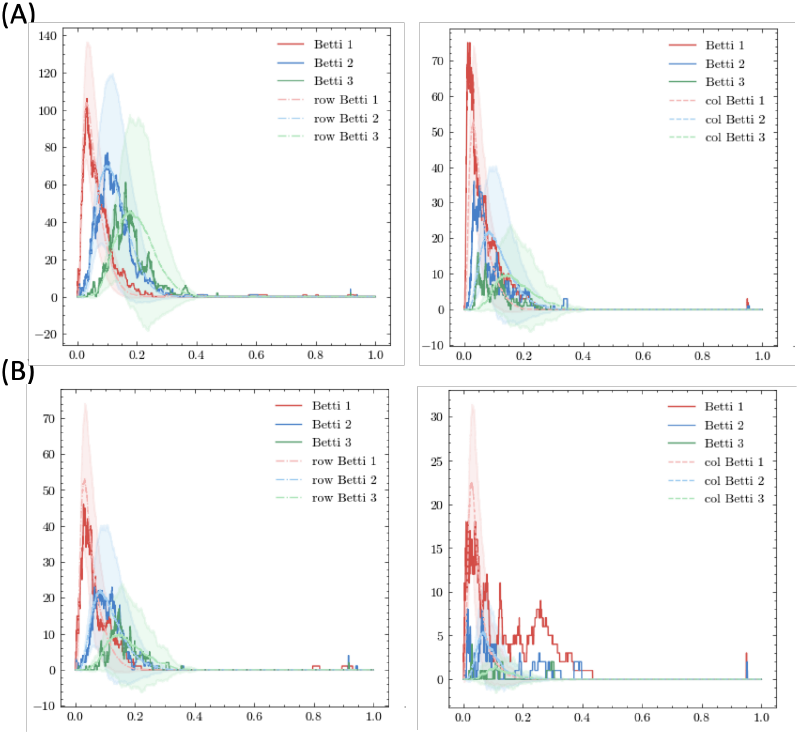
2D Hyperbolic Embedding of the Structural Representation Among Proofread Excitatory Neurons in 4P: Similar to Figure S12, this analysis focuses on layer 4 pyramidal cells (*N* = 272) using (A) binary connectome or (B) weighted connectome to evaluate **C**^out^ (left column) and **C**^in^ (right column). Synthetic data are presented with optimized *R*_max_, averaged across 100 trials.

**Fig. S15.**
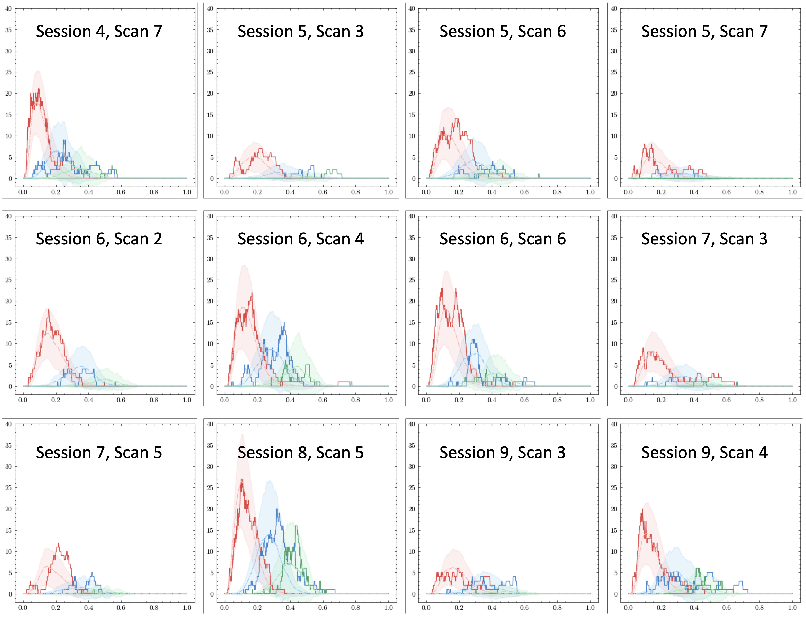
Betti Analysis of All Activity Traces Indicates a Universal Hyperbolic Embedding: Repeat the analysis in Figure 1E for all scans by comparing the alignment of the top three Betti curves between groundtruth and synthetic data, averaged across 100 trials. Use the parameters described in Section 4.2, with varying *R*_max_ values optimized within the range of 5 to 20 (grid size = 0.1).

**Fig. S16.**
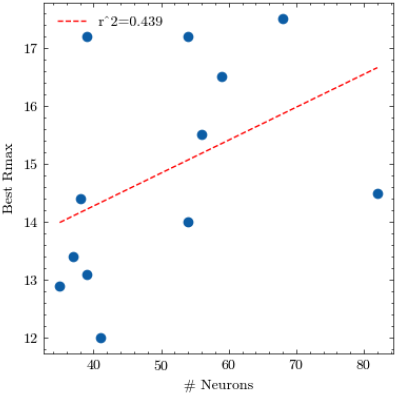
Positive Relationship Between the Number of Neurons in the Scan and the Embedded Hyperbolic Radius: Results derived from Figure S15; including more neurons increase the effective dimensionality of the representation space, thereby necessitating greater capacity.

**Fig. S17.**
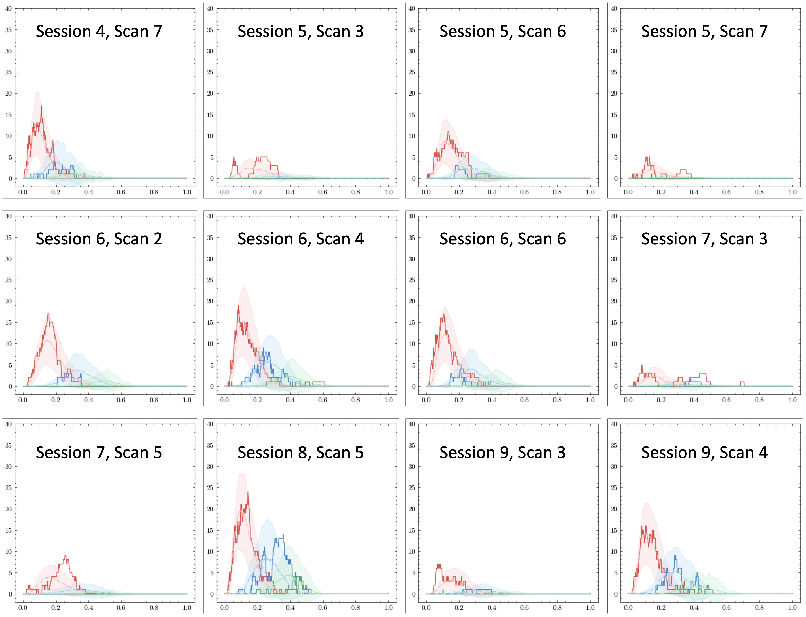
Betti Analysis of All Activity Traces Using Cosine Similarity: Similar to Figure S15, but using cosine similarity to quantify activity proximity.

**Fig. S18.**
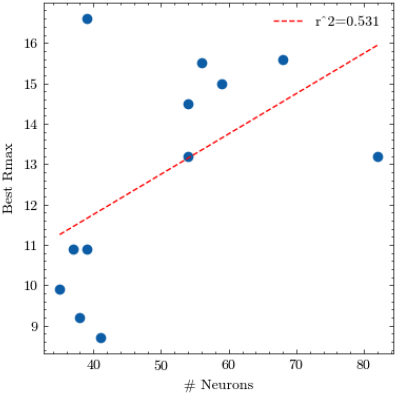
Positive Relationship Between the Number of Neurons in the Scan and the Embedded Hyperbolic Radius Using Cosine Similarity as the Correlation Measure: Results derived from Figure S17; Similar to Figure S16. Under this measure, the representation space becomes less complex, reducing the magnitude of the Betti curves and an overall decrease in the hyperbolic radii *R*_max_.

**Fig. S19.**
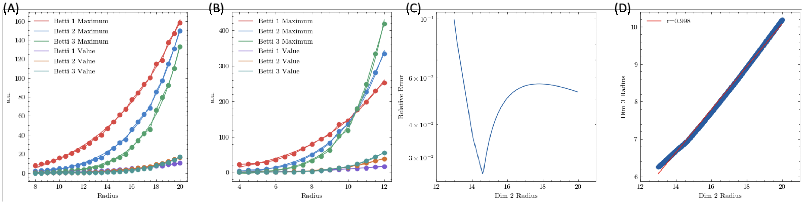
Transformation between 2D and 3D hyperbolic embedding can be achieved through an approximately linear relationship of *R*_max_: Specific case for *N* = 217 and noise level *ϵ* = 0.0625 averaged over 100 trials. (A-B): Presentations of 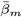 and max(*β*_*m*_(*ρ*)), *m* = 1, 2, 3 for 2D (A) and 3D (B). Each curve is interpolated using a simple exponential function and transformed into the continuous case; (C-D): Optimized cumulative pointwise error (C) and the mapping relationship (D) between 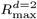 and 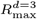. Note that when 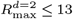, the 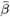 values are small (B part), which could lead to stability issues.

**Table S5.** Performance (*acc*) Comparison between Reconstructing Raw Fluorescence Traces and Detrended, Deconvolved Traces (Used in Main Text) via the Linear Model: Different forms of **C** are used to compute the coefficients ***α*** under the baseline experiment setup (Section 2.3 and Figure 3).

|  | Raw Trace | Deconvolved Trace |
| --- | --- | --- |
| HypEmbed $C^{\text{in}}$ | 24.1% | 8.66% |
| $C^{\text{in}}$ | 23.1% | 7.70% |
| HypEmbed $C^{\text{out}}$ | 22.8% | 8.05% |
| $C^{\text{out}}$ | 15.8% | 4.92% |
| $C^{\text{act}}$ | 36.4% | 13.4% |
| Mean of $C^{\text{act}}$ | 7.00% | 1.36% |

**Fig. S20.**
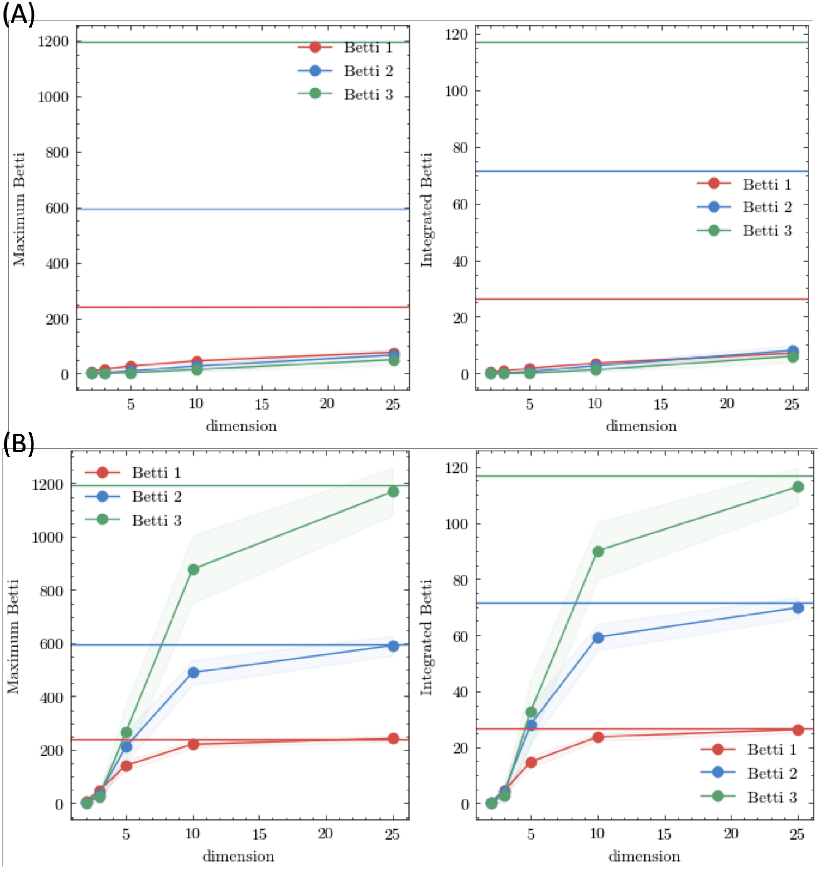
Comparison of Random Matrices and Distances Matrices Sampled from Hyperbolic and Euclidean Spaces Across Various Dimensions: Variations in Betti values 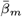 (left column) and maximum Betti curve value max(*β*_*m*_(*ρ*)) (right column) for *m* = 1, 2, 3 are presented across different dimensions with *N* = 100 for (A) Euclidean space and (B) hyperbolic space, using the parameters specified in Section 4.2. Horizontal lines indicate benchmark outcomes derived from random matrices. Each experiment is repeated for 500 trials.

**Fig. S21.**
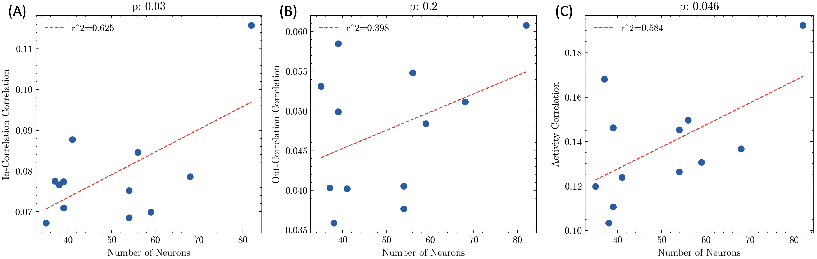
Having More Functionally Coregistered Neurons in a Single Scan Enhances Reconstruction Accuracy: This result remains consistent when using (A) **C**^in^, (B): **C**^out^, and (C): **C**^act^. This example considers only proofread neurons with the weighted connectome, but the result is applicable to various settings.

**Fig. S22.**
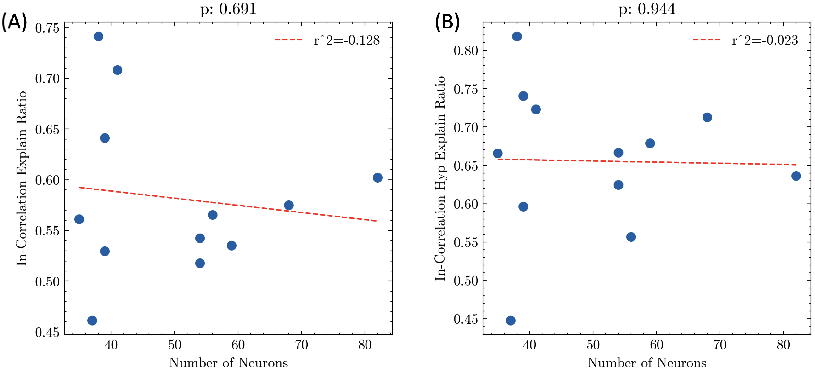
Having More Functionally Coregistered Neurons in a Single Scan Does Not Improve the Explanation Ratio, Regardless of Full Connectome Information or Embedded Representation: Compared to Figure S21, although the absolute reconstruction accuracy *a* significantly improves with increasing *N*, the explanation ratio (*a/a*_act_ ) remains uncorrelated with *N* for both the full connectome **C**^in^ (left) and its hyperbolic embeddings (right). The observation holds consistently for output correlation (not shown). Data points correspond to the green and red parts in Figure 3.

**Fig. S23.**
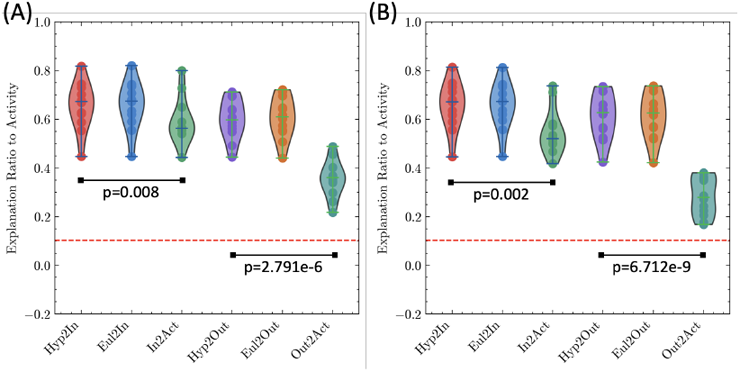
**Calculate C^in^ and C^out^ (and embeddings) using (A) binary connectome (A) or (B) postsynaptic density (PSD) information**.

**Fig. S24.**
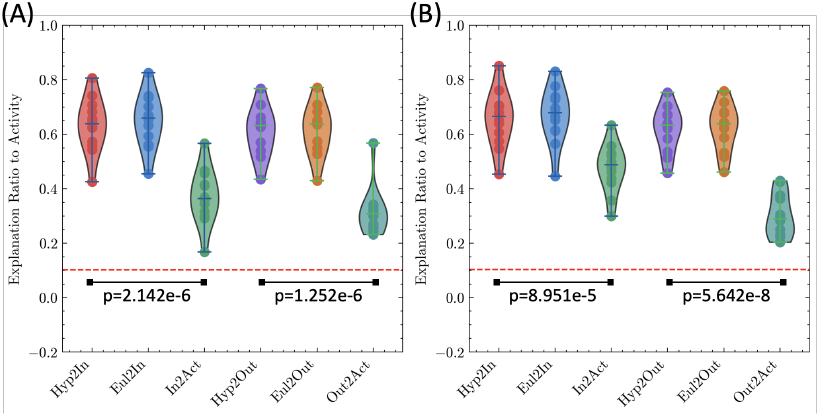
**Calculate C^in^ and C^out^ (and embeddings) using weighted connectome to/from only (A) excitatory neurons and (B) inhibitory neurons**.

**Fig. S25.**
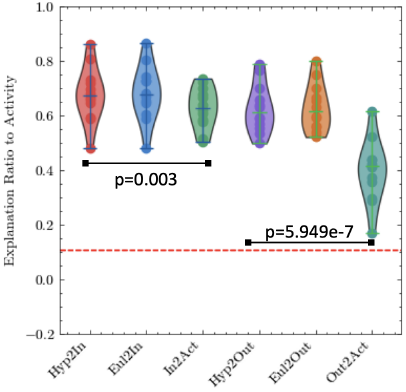
**Calculate C^in^ and C^out^ (and embeddings) using structural information from all excitatory and inhibitory neurons, including not fully proofread ones.**

**Fig. S26.**
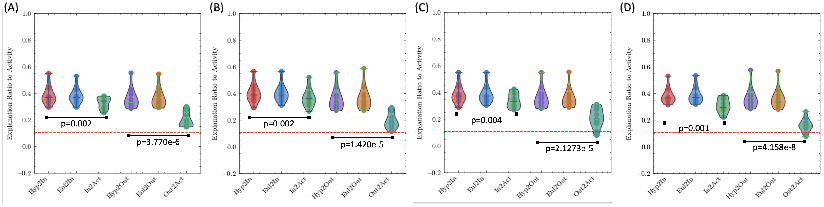
**Calculate C^act^ respectively for each visual stimulus period based on (A) synaptic count with proofread neurons, (B) synaptic count with all E/I neurons, (C) binary connectome with proofread neurons, and (D) PSD with proofread neurons to calculate structural similarity.**

**Fig. S27.**
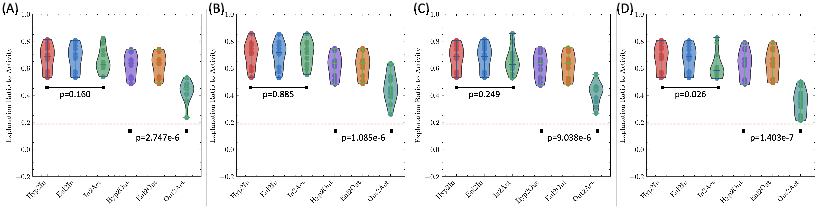
Reconstruction of Raw Fluorescence Traces based on (A) synaptic count with proofread neurons, (B) synaptic count with all E/I neurons, (C) binary connectome with proofread neurons, and (D) PSD sizes with proofread neurons to calculate structural similarity: The explanation ratio based on the median pairwise neuronal activity increases to 18.6% (Figure 3).

**Fig. S28.**
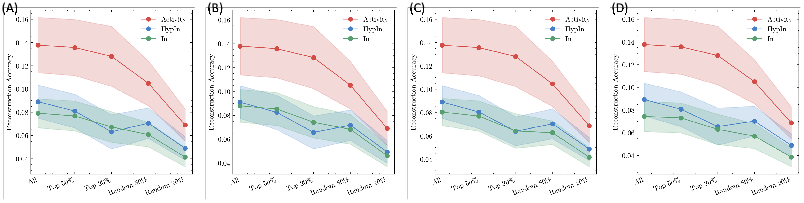
**Reconstruction using Partially or Randomly Selected Neuronal Activity Traces (averaged over 100 trials), based on (A) synaptic count with proofread neurons, (B) synaptic count with all E/I neurons, (C) binary connectome with proofread neurons, and (D) PSD sizes with proofread neurons to compute structural similarity. Means and standard deviations are calculated across 12 independent scans.**

**Fig. S29.**
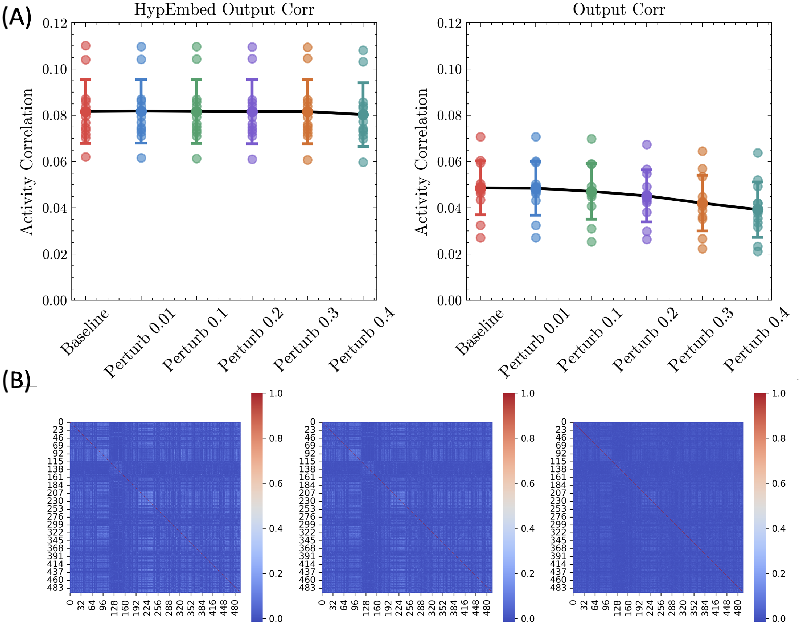
Comparison of Reconstruction Accuracy Using Hyperbolic Embedding of Output Similarity versus Output Similarity: (A): Reconstruction accuracy of the hyperbolic embedding of **C**^out^ (left) and **C**^out^ itself (right) under the baseline experiment (Figure 3) and varying levels of perturbation. (B) Illustration of **C**^out^ among targeted neurons (*N* = 497) for the baseline experiment (left), 10% perturbation (middle), and 40% perturbation (right). In (A), each scatted point represents the average result across 10 random perturbations for an individual scan. The mean and standard deviation across all scans are also shown. In (B), higher perturbation diminishes the grouping structure of neurons, resulting in a more homogeneous outcome.

**Fig. S30.**
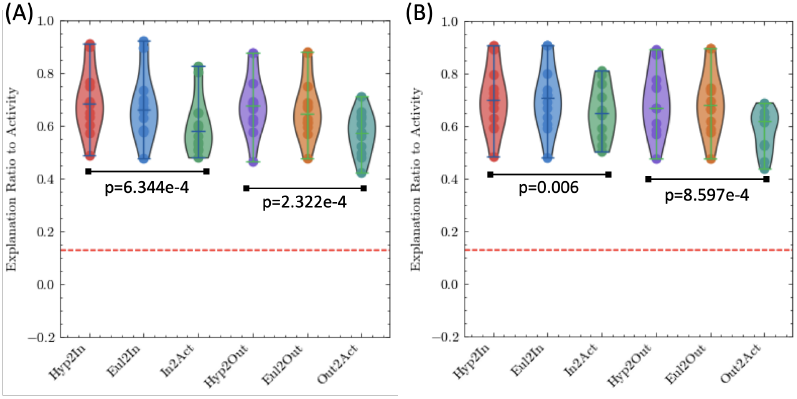
Reconstruction using activity from proofread neurons based on (A) synaptic count with proofread neurons and (B) synaptic count with all E/I neurons to calculate structural similarity: The graph illustrates explanation ratio (*a*/*a*_act_). As discussed in Section 2.3 and Figure S21, overall accuracies decrease (except the output reliability) due to the stricter criterion and less neurons included per scan. For (A), **C**^in^ achieves 6.78% and **C**^out^ achieves 6.50%, while the corresponding counterpart values are 7.17% and 4.65%, respectively. (B) has 7.28% and 6.65%, and the counterpart has 8.08% and 5.01%.

**Fig. S31.**
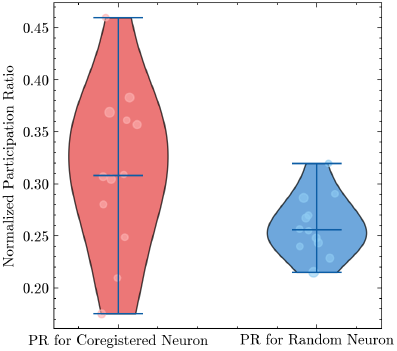
Dimensionality of Activity Space: In the main text, we exclusively considered neurons that are functionally coregistered (i.e. neurons for which connectome information, though potentially incomplete, is available). We quantify the dimensionality of the activity space formed by these *N*_*i*_ neurons using the normalized participation ratio (NPR) (58). Additionally, we evaluate the NPR for randomly selected sets of *N*_*i*_ neurons in each scan, averaged over 100 instances. The size of the scattered points is scaled with *N*_*i*_.

**Fig. S32.**
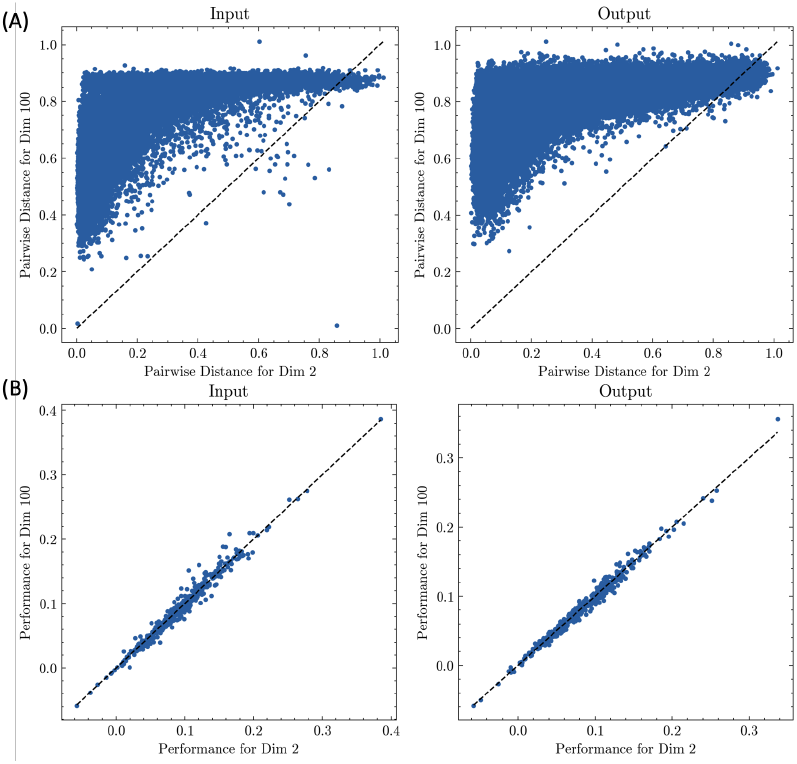
Comparison of (A) pairwise distances between Euclidean embedded points and (B) neuron-wise reconstruction accuracy, concatenated across all scans, for different dimensionality: We compare the results using optimized Euclidean embeddings in ℝ^2^ and ℝ^100^. Despite significant variation in distance preservation quality, the neuron-wise reconstruction accuracy remains consistent. In (B), we averaged the performance (Pearson correlation) for neurons included in multiple scans. Compared to Section 4.1, ℝ^100^ would still be considered “low-dimensional” given the dataset size. Although it achieves near zero stress (not shown), the rank preservation (of pairwise distance remains imperfect (*R*^2^ = 0.73 for the input and *R*^2^ = 0.92 for the output), highlighting its distinction from the raw connectome data. Due to the computational challenges and instability caused by the high degree of freedom in “high”-dimensional embeddings, we limit the test to ℝ^100^.

**Fig. S33.**
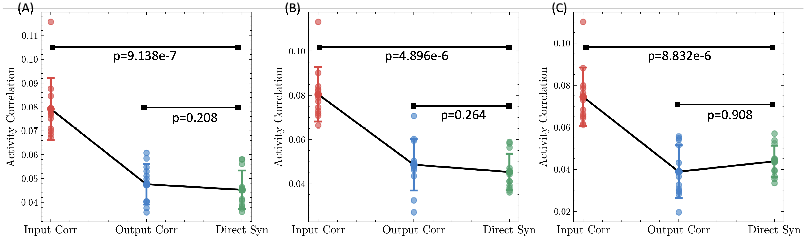
Comparison Between Using Normalized Direct Connectivity as Coefficients in the Linear Model and Using Input and Output Similarity: We compare the reconstruction performance in Figure 3 and Figure S23, using input and output correlation respectively versus direct connectivity measures. The direct connectivity is represented through (A) synaptic count, (B) binary connectivity, and (C) PSD sizes. Means and standard deviations are reported.

**Fig. S34.**
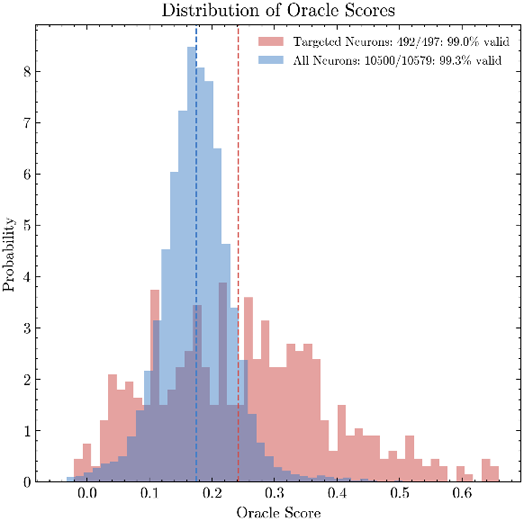
Distribution of Oracle Scores for Targeted Neurons (*N* = 497) and All Functionally Coregistered Neurons (*N* = 10579): Only values not reported as `NaN` are shown (7). For neurons included in multiple scans, we report the mean oracle score. The median value for targeted neurons (0.242) is slightly higher than that of the overall population (0.175).

**Fig. S35.**
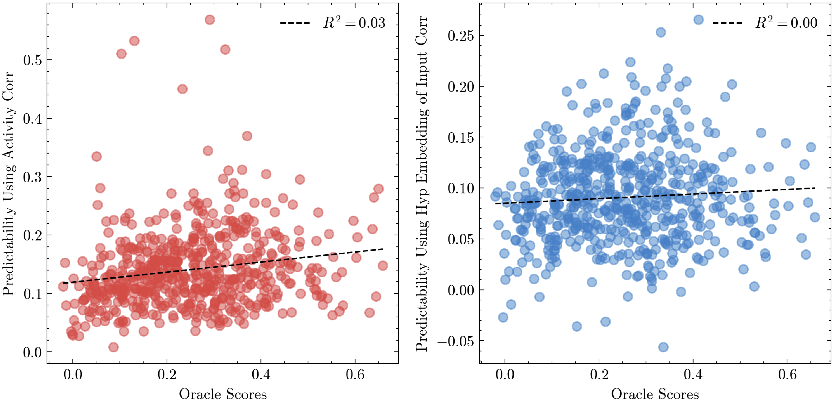
Comparison between Oracle Score and Neuron-wise Predictability: (A): Correlation between oracle score and *acc*_act_, obtained by using **C**^act^ in the baseline model to set the weights of the linear coefficients ***α*** (Figure 2 and 3); (B): Correlation with **C**^hyp,in^. The *R*^2^ value from linear regression is reported in the legend. All occurrences of targeted neurons with valid oracle score (including duplicates) across the 12 scans (*N* = 585) are included (Figure S34). These results indicate that neuron-wise predictability, as defined by our linear model, cannot be simply explained by the oracle score (7).

## Footnotes

1 https://github.com/StevenZhang0116/StructureFunctionAlignment

2 https://github.com/nerdnik/PyCliqueTop_2023

3 https://github.com/gyrheart/Hyperbolic-MDS

4 https://github.com/gyrheart/Hyperbolic-t-SNE

